# DeepAD: Alzheimer’s Disease Classification via Deep Convolutional Neural Networks using MRI and fMRI

**DOI:** 10.1101/070441

**Authors:** Saman Sarraf, Danielle D. DeSouza, John Anderson, Ghassem Tofighi, for the Alzheimer's Disease Neuroimaging Initiativ

## Abstract

To extract patterns from neuroimaging data, various statistical methods and machine learning algorithms have been explored for the diagnosis of Alzheimer’s disease among older adults in both clinical and research applications; however, distinguishing between Alzheimer’s and healthy brain data has been challenging in older adults (age > 75) due to highly similar patterns of brain atrophy and image intensities. Recently, cutting-edge deep learning technologies have rapidly expanded into numerous fields, including medical image analysis. This paper outlines state-of-the-art deep learning-based pipelines employed to distinguish Alzheimer’s magnetic resonance imaging (MRI) and functional MRI (fMRI) from normal healthy control data for a given age group. Using these pipelines, which were executed on a GPU-based high-performance computing platform, the data were strictly and carefully preprocessed. Next, scale- and shift-invariant low- to high-level features were obtained from a high volume of training images using convolutional neural network (CNN) architecture. In this study, fMRI data were used for the first time in deep learning applications for the purposes of medical image analysis and Alzheimer’s disease prediction. These proposed and implemented pipelines, which demonstrate a significant improvement in classification output over other studies, resulted in high and reproducible accuracy rates of 99.9% and 98.84% for the fMRI and MRI pipelines, respectively. Additionally, for clinical purposes, subject-level classification was performed, resulting in an average accuracy rate of 94.32% and 97.88% for the fMRI and MRI pipelines, respectively. Finally, a decision making algorithm designed for the subject-level classification improved the rate to 97.77% for fMRI and 100% for MRI pipelines.

## 2 Introduction

### 2.1 Alzheimer’s Disease

Alzheimer’s disease (AD) is an irreversible, progressive neurological brain disorder and multifaceted disease that slowly destroys brain cells, causing memory and thinking skill losses, and ultimately loss of the ability to carry out even the simplest tasks. The cognitive decline caused by this disorder ultimately leads to dementia. For instance, the disease begins with mild deterioration and grows progressively worse as a neurodegenerative type of dementia. Diagnosing Alzheimer’s disease requires very careful medical assessment, including patient history, a mini mental state examination (MMSE), and physical and neurobiological exams [1] [2]. In addition to these evaluations, structural magnetic resonance imaging and resting state functional magnetic resonance imaging (rs-fMRI) offer non-invasive methods of studying the structure of the brain, functional brain activity, and changes in the brain. During scanning using both structural (anatomical) and rs-fMRI techniques, patients remain prone on the MRI table and do not perform any tasks. This allows data acquisition to occur without any effects from a particular task on functional activity in the brain [3] [4] [5]. Alzheimer’s disease causes shrinkage of the hippocampus and cerebral cortex and enlargement of ventricles in the brain. The level of these effects is dependent upon the stage of disease progression. In the advanced stage of AD, severe shrinkage of the hippocampus and cerebral cortex, as well as significantly enlarged ventricles, can easily be recognized in MR images. This damage affects those brain regions and networks related to thinking, remembering (especially short-term memory), planning and judgment. Since brain cells in the damaged regions have degenerated, MR image (or signal) intensities are low in both MRI and rs-fMRI techniques [6] [7] [8]. However, some of the signs found in the AD imaging data are also identified in normal aging imaging data. Identifying the visual distinction between AD data and images of older subjects with normal aging effects requires extensive knowledge and experience, which must then be combined with additional clinical results in order to accurately classify the data (i.e., MMSE) [1]. Development of an assistive tool or algorithm to classify MR-based imaging data, such as structural MRI and rs-fMRI data, and, more importantly, to distinguish brain disorder data from healthy subjects, has always been of interest to clinicians [9]. A robust machine learning algorithm such as Deep Learning, which is able to classify Alzheimer’s disease, will assist scientists and clinicians in diagnosing this brain disorder and will also aid in the accurate and timely diagnosis of Alzheimer’s patients [10].

### 2.2 Deep Learning

Hierarchical or structured deep learning is a modern branch of machine learning that was inspired by the human brain. This technique has been developed based upon complicated algorithms that model high-level features and extract those abstractions from data by using similar neural network architecture that is actually much more complicated. Neuroscientists have discovered that the “neocortex,” which is a part of the cerebral cortex concerned with sight and hearing in mammals, processes sensory signals by propagating them through a complex hierarchy over time. This served as the primary motivation for the development of deep machine learning that focuses on computational models for information representation which exhibits characteristics similar to those of the neocortex [11] [12] [13].

Convolutional neural networks (CNNs) that are inspired by the human visual system are similar to classic neural networks. This architecture has been specifically designed based on the explicit assumption that raw data are comprised of two-dimensional images that enable certain properties to be encoded while also reducing the amount of hyper parameters. The topology of CNNs utilizes spatial relationships to reduce the number of parameters that must be learned, thus improving upon general feed-forward backpropagation training [14] [15]. Equation 1 demonstrates how the gradient component for a given weight is calculated in the backpropagation step, where E is error function, y is the neuron N_i,j_, x is the input, l represents layer numbers, w is filter weight with a and b indices, N is the number of neurons in a given layer, and m is the filter size.

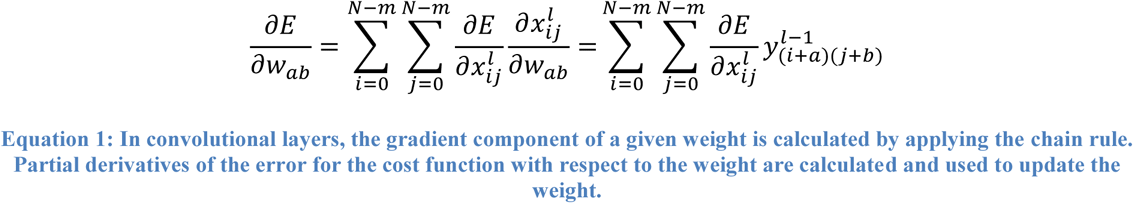

Equation 2 describes the backpropagation error for the previous layer using the chain rule. This equation is similar to the convolution definition, where *x*_(*i+a*)(*j+b*)_ is replaced by *x*_(*i*−*a*)(*j*−*b*)_. It demonstrates the backpropagation results in convolution while the weights are rotated. The rotation of the weights derives from a delta error in the convolutional neural network.

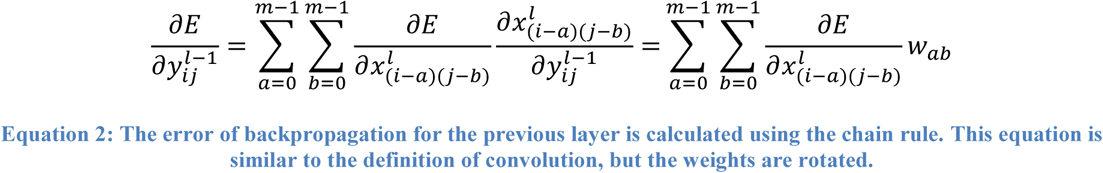

In CNNs, small portions of the image (called local receptive fields) are treated as inputs to the lowest layer of the hierarchical structure. One of the most important features of CNNs is that their complex architecture provides a level of invariance to shift, scale and rotation, as the local receptive field allows the neurons or processing units access to elementary features, such as oriented edges or corners. This network is primarily comprised of neurons having learnable weights and biases, forming the convolutional layer. It also includes other network structures, such as a pooling layer, a normalization layer and a fully connected layer. As briefly mentioned above, the convolutional layer, or conv layer, computes the output of neurons that are connected to local regions in the input, each computing a dot product between its weight and the region it is connected to in the input volume. The pooling layer, also known as the pool layer, performs a downsampling operation along the spatial dimensions. The normalization layer, also known as the rectified linear units (ReLU) layer, applies an elementwise activation function, such as max (0, x) thresholding at zero. This layer does not change the size of the image volume [11] [12] [16]. The fully connected (FC) layer computes the class scores, resulting in the volume of the number of classes. As with ordinary neural networks, and as the name implies, each neuron in this layer is connected to all of the numbers in the previous volume [12] [17]. The convolutional layer plays an important role in CNN architecture and is the core building block in this network. The conv layer’s parameters consist of a set of learnable filters. Every filter is spatially small but extends through the full depth of the input volume. During the forward pass, each filter is convolved across the width and height of the input volume, producing a 2D activation map of that filter. During this convolving, the network learns of filters that activate when they see some specific type of feature at some spatial position in the input. Next, these activation maps are stacked for all filters along the depth dimension, which forms the full output volume. Every entry in the output volume can thus also be interpreted as an output from a neuron that only examines a small region in the input and shares parameters with neurons in the same activation map [12] [18]. A pooling layer is usually inserted between successive conv layers in CNN architecture. Its function is to reduce (down sample) the spatial size of the representation in order to minimize network hyper parameters, and hence also to control overfitting. The pooling layer operates independently on every depth slice of the input and resizes it spatially using the max operation [11] [12] [16] [17] [18]. In convolutional neural network architecture, the conv layer can accept any image (volume) of size *W*_1_*×H*_1_*× D*_1_ that also requires four hyper parameters, which are *K*, number of filters; *F*, their spatial extent; *S*, the size of stride; and *P,* the amount of zero padding. The conv layer outputs the new image, whose dimensions are *W*_2_ × *H*_2_ × *D*_2_, calculated as Equation 3 below:

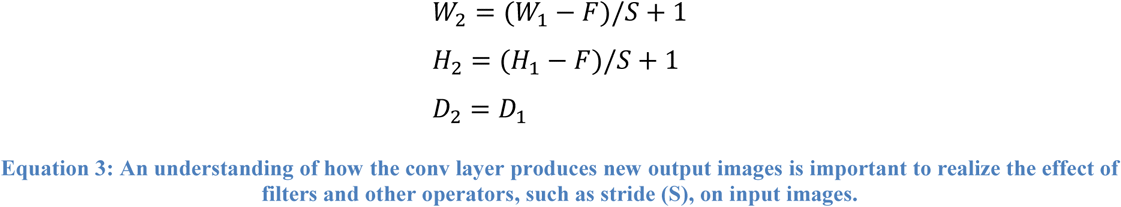

LeNet-5 was first designed by Y. LeCun et al. [11]. This architecture successfully classified digits and was applied to hand-written check numbers. The application of this fundamental but deep network architecture expanded into more complicated problems by adjusting the network hyper parameters.

LeNet-5 architecture, which extracts low- to mid-level features, includes two conv layers, two pooling layers, and two fully connected layers, as shown in Figure 1.

**Figure 1.**
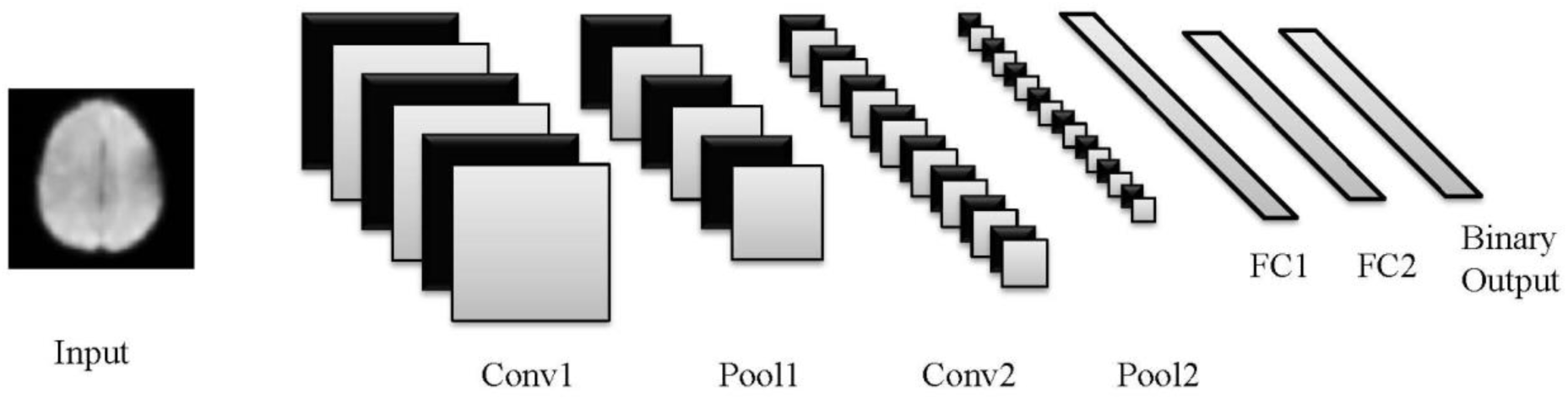
**LeNet-5 includes two conv, two pool and two FC layers. The original version of this network classified 10 digits. In this work, the architecture was optimized for binary output, which were Alzheimer’s disease (AD) and normal control (NC), respectively.**

More complex CNN architecture was developed to recognize numerous objects derived from high volume data, including AlexNet (ImageNet) [19], ZF Net [20], GoogleNet [17], VGGNet [21] and ResNet [22]. GoogleNet, which was developed by Szegedy et al. [17], is a successful network that is broadly used for object recognition and classification. This architecture is comprised of a deep, 22-layer network based on a modern design module called Inception. One of the fundamental approaches to improving the accuracy of CNN architecture is to increase the size of layers. However, this straightforward solution causes two major issues. First, a large number of hyper parameters requires more training data and may also result in overfitting, especially in the case of limited training data. On the other hand, uniform increases in network size dramatically increase interactions with computational resources, which affect the timing performance and the cost of providing infrastructure. One of the optimized solutions for both problems would be the development of a sparsely connected architecture rather than a fully connected network. Strict mathematical proofs demonstrate that the well-known Hebbian principle of neurons firing and wiring together created the Inception architecture of GoogleNet [17]. The Inception module of GoogleNet, as shown in Figure 2, is developed by discovering the optimal local sparse structure to construct convolutional blocks. Inception architecture allows for a significant increase in the number of units at each layer, while computational complexity remains under control at later stages, which is achieved through global dimensionality reduction prior to costly convolutions with larger patch sizes.

**Figure 2.**
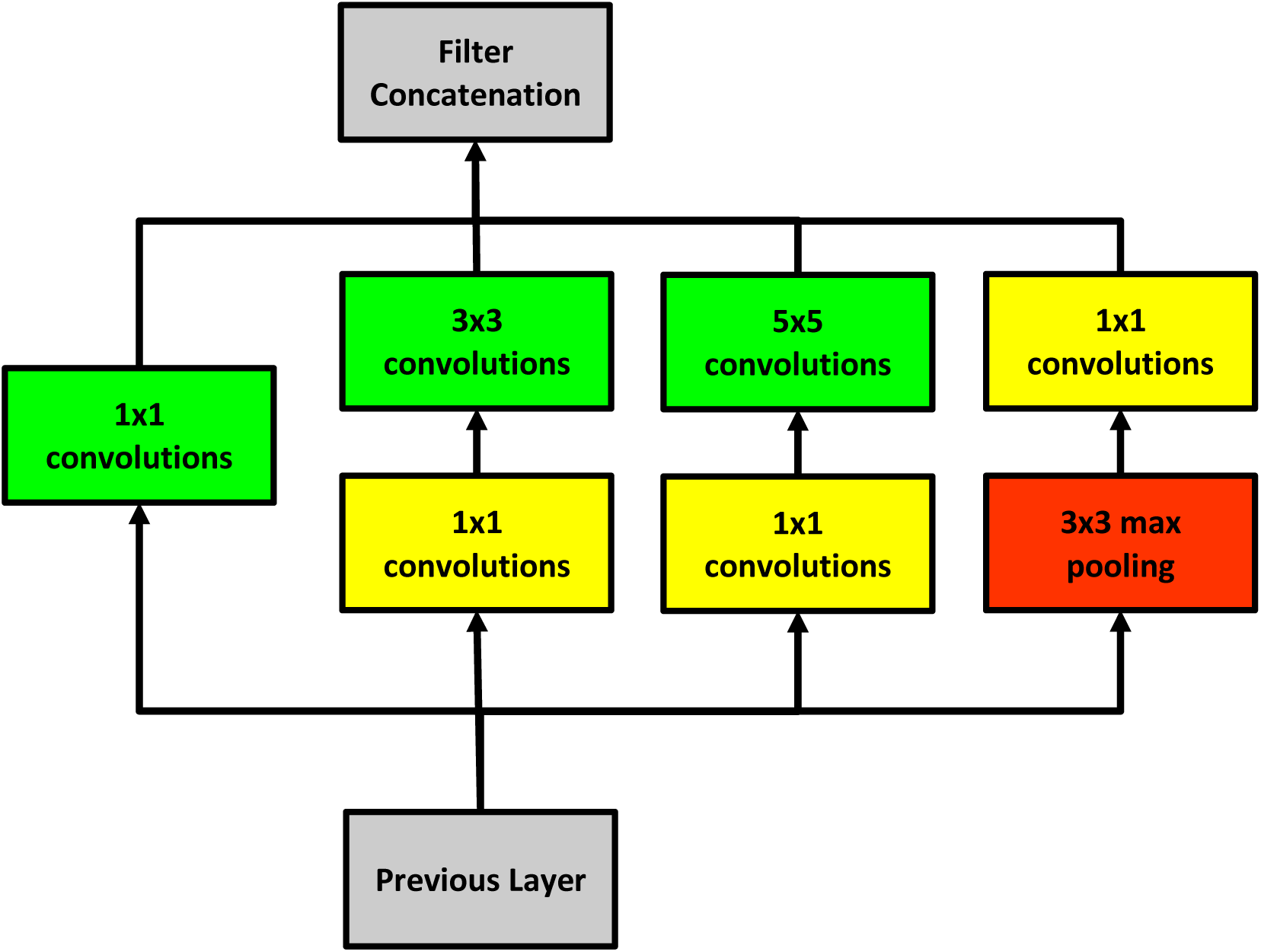
**Inception module with dimensionality reduction in GoogleNet architecture.**

### 2.3 Data Acquisition

In this study, two subsets of the ADNI database (http://adni.loni.usc.edu/) were used to train and validate convolutional neural network classifiers. The first subset included 144 subjects who were scanned for resting-state functional magnetic resonance imaging (rs-fMRI) studies. In this dataset, 52 Alzheimer’s patients and 92 healthy control subjects were recruited (age group > 75). The second dataset included 302 subjects whose structural magnetic resonance imaging data (MRI) were acquired (age group > 75). This group included 211 Alzheimer’s patients and 91 healthy control subjects. Certain subjects were scanned at different points in time, and their imaging data were separately considered in this work. Table 1 presents the demographic information for both subsets, including mini mental state examination (MMSE) scores.

**Table 1.** **Two subsets of the ADNI database were used in this study, including 144 subjects with fMRI data and 302 subjects with MRI data. The mean and standard deviation (SD) of age and total MMSE scores per group are delineated in the table below.**

| Modality | Total Subj. | Group | Subj. | Female | Mean of Age | SD | Male | Mean of Age | SD | MMSE ± SD |
| --- | --- | --- | --- | --- | --- | --- | --- | --- | --- | --- |
| rs-fMRI | 144 | Alzheimer | 52 | 21 | 79.42 | 16.35 | 31 | 80.54 | 15.98 | 22.70 ± 2.10 |
| | | Control | 92 | 43 | 80.79 | 19.16 | 49 | 81.75 | 21.43 | $28.82 \pm 1.35$ |
| MRI | 302 | Alzheimer | 211 | 85 | 80.98 | 21.6 | 126 | 81.27 | 16.66 | $23.07 \pm 2.06$ |
| | | Control | 91 | 43 | 79.37 | 12.52 | 48 | 80.81 | 19.51 | $28.81 \pm 1.35$ |

MRI data acquisition was performed according to the ADNI acquisition protocol [23]. Scanning was performed on three different Tesla scanners, General Electric (GE) Healthcare, Philips Medical Systems, and Siemens Medical Solutions, and was based on identical scanning parameters. Anatomical scans were acquired with a 3D MPRAGE sequence (TR=2s, TE=2.63 ms, FOV=25.6 cm, 256 × 256 matrix, 160 slices of 1mm thickness). Functional scans were acquired using an EPI sequence (150 volumes, TR=2 s, TE=30 ms, flip angle=70, FOV=20 cm, 64 × 64 matrix, 30 axial slices of 5mm thickness without gap).

## 3 Related Work

Changes in brain structure and function caused by Alzheimer’s disease have proved of great interest to numerous scientists and research groups. In diagnostic imaging in particular, classification and predictive modeling of the stages of Alzheimer’s have been broadly investigated. Suk et al. [24] [25] [26] developed a deep learning-based method to classify AD magnetic current imaging (MCI) and MCI-converter structural MRI and PET data, achieving accuracy rates of 95.9%, 85.0% and 75.8% for the mentioned classes, respectively. In their approach, Suk et al. developed an auto-encoder network to extract low- to mid-level features from images. Next, classification was performed using multi-task and multi-kernel Support Vector Machine (SVM) learning methods. This pipeline was improved by using more complicated SVM kernels and multimodal MRI/PET data. However, the best accuracy rate for Suk et al. remained unchanged [27]. Payan et al. [28] of Imperial College London designed a predictive algorithm to distinguish AD MCI from normal healthy control subjects’ imaging. In this study, an auto-encoder with 3D convolutional neural network architecture was utilized. Payan et al. obtained an accuracy rate of 95.39% in distinguishing AD from NC subjects. The research group also tested a 2D CNN architecture, and the reported accuracy rate was nearly identical in terms of value. Additionally, a multimodal neuroimaging feature extraction pipeline for multiclass AD diagnosis was developed by Liu et al. [29]. This deep-learning framework was developed using a zero-masking strategy to preserve all possible information encoded in imaging data. High-level features were extracted using stacked auto-encoder (SAE) networks, and classification was performed using SVM against multimodal and multiclass MR/PET data. The highest accuracy rate achieved in that study was 86.86%. Aversen et al. [30], Liu et al. [31], Siqi et al. [32], Brosch et al. [33], Rampasek et al. [34], De Brebisson et al. [35] and Ijjina et al. [36] also demonstrated the application of deep learning in automatic classification of Alzheimer’s disease from structural MRI, where AD, MCI and NC data were classified.

**Table 2.** **The table below summarizes the related works and percentage accuracies reported in each reference. Using MRI and PET data have been interest of researchers and different techniques were utilized to classify Alzheimer’s disease from Normal Control (NC) or Mild Cognitive Impairment (MCI). As seen, resting-state functional MRI (rs-fMRI) and a full deep learning-based pipeline were used as a novel approach. Also, the end to end pipeline improved the accuracy rate for structural MRI data. The abbreviations in the table are as following: CAE: Convolutional Autoencoder, SAE: Sparse Autoencoder, CNN: Convolutional Neural Networks, MSLF: Matrix-Similarity based Loss Function, MTFS: Multi-task feature selection, DL: Deep Learning and MFE: Multiview Feature Extraction. *: slice level classification, **: subject level classification followed by the decision making algorithm.**

| Reference | Modality | Method | AD/MCI/NC | AD+MCI/NC | AD/NC | AD/MCI | NC/MCI |
| --- | --- | --- | --- | --- | --- | --- | --- |
| Suk et al. [24] | PET+MRI+CSF | SAE+SVM | N/A | N/A | 95.9 | N/A | 85 |
| Suk et al. [27] | PET+MRI | SAE+SVM | N/A | N/A | 95.4 | N/A | 85.7 |
| Zhu et al. [37] | PET+MRI+CSF | MSLF+SVM | N/A | N/A | 95.9 | N/A | 82 |
| Zu et al. [38] | PET+MRI | MTFS+SVM | N/A | N/A | 96 | N/A | 80.3 |
| Liu et al. [39] | PET+MRI | SAE+SVM | 53.8 | N/A | 91.4 | N/A | 82.1 |
| Liu et al. [40] | MRI | MFE+SVM | N/A | N/A | 93.8 | N/A | 89.1 |
| Li et al. [41] | PET+MRI+CSF | PCA+SVM | N/A | N/A | 91.4 | 70.1 | 77.4 |
| Payan et al. [28] | MRI | 2D-SAE | 89.4 | N/A | 95.4 | 86.8 | 92.1 |
| Hosseini-Asl [42] | MRI | 3D-CAE | 89.1 | 90.3 | 97.6 | N/A | N/A |
| Sarraf et al. | rs-fMRI* | DL-CNN | N/A | N/A | 99.9 | N/A | N/A |
| Sarraf et al. | MRI* | DL-CNN | N/A | N/A | 98.84 | N/A | N/A |
| Sarraf et al. | rs-fMRI** | DL-CNN | N/A | N/A | 97.77 | N/A | N/A |
| Sarraf et al. | MRI** | DL-CNN | N/A | N/A | 100 | N/A | N/A |

## 4 Methods

Classification of Alzheimer’s disease images and normal, healthy images required several steps, from preprocessing to recognition, which resulted in the development of an end-to-end pipeline. Three major modules formed this recognition pipeline: a) preprocessing b) data conversion; and c) classification, respectively. Two different approaches were used in the preprocessing module, as preprocessing of 4D rs-fMRI and 3D structural MRI data required different methodologies, which will be explained later in this paper. After the preprocessing steps, the data were converted from medical imaging to a lossless Portable Network Graphics (PNG) format to input into the deep learning-based classifier. Finally, the CNN-based architecture receiving images in its input layer was trained and tested (validated) using 75% and 25% of the dataset, respectively. In practice, two different pipelines were developed, each of which was different in terms of preprocessing but similar in terms of data conversion and classification steps, as demonstrated in Figure 3.

**Figure 3.**
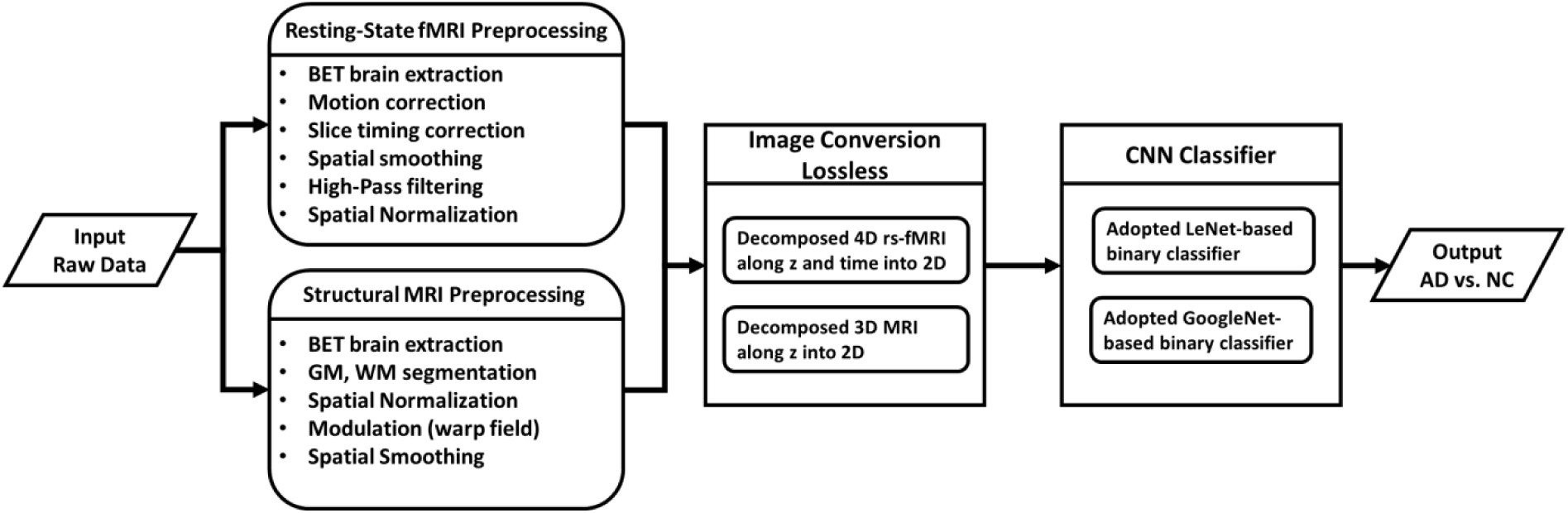
**End-to-end recognition based on deep learning CNN classification methods is comprised of three major components: preprocessing, image conversion and classification modules. In the preprocessing step, two different submodules were developed for rs-fMRI and structural data. Next, the lossless image conversion module created PNG images from medical imaging data using the algorithm described in the following section of this paper. The final step was to recognize AD from NC samples using CNN models, which was performed by training and testing models using 75% and 25% of the samples, respectively.**

### 4.1 rs-fMRI Data Preprocessing

The raw data in DICOM format for both the Alzheimer’s (AD) group and the normal control (NC) group were converted to NII format (Neuroimaging Informatics Technology Initiative - NIfTI) using the dcm2nii software package developed by Chris Roden et al. http://www.sph.sc.edu/comd/rorden/mricron/dcm2nii.html. Next, non-brain regions, including skull and neck voxels, were removed from the structural T1-weighted image corresponding to each fMRI time course using FSL-BET [43]. Resting-state fMRI data, including 140 time series per subject, were corrected for motion artefact using FSL-MCFLIRT [44], as low frequency drifts and motion could adversely affect decomposition. The next necessary step was the regular slice timing correction, which was applied to each voxel’s time series because of the assumption that later processing assumes all slices were acquired exactly half-way through the relevant volume’s acquisition time (TR). In fact, each slice is taken at slightly different times. Slice timing correction works by using Hanning-windowed Sinc interpolation to shift each time series by an appropriate fraction of a TR relative to the middle of the TR period. Spatial smoothing of each functional time course was then performed using a Gaussian kernel of 5 mm full width at half maximum. Additionally, low-level noise was removed from the data by a temporal high-pass filter with a cut-off frequency of 0.01 HZ (sigma = 90 seconds) in order to control the longest allowed temporal period. The functional images were registered to the individual’s high-resolution (structural T1) scan using affine linear transformation with seven degrees of freedom (7 DOF). Subsequently, the registered images were aligned to the MNI152 standard space (average T1 brain image constructed from 152 normal subjects at the Montreal Neurological Institute) using affine linear registration with 12 DOF followed by 4 mm resampling, which resulted in 45×54×45 images per time course.

### 4.2 Structural MRI Data Preprocessing

The raw data of structural MRI scans for both the AD and the NC groups were provided in NII format in the ADNI database. First, all non-brain tissues were removed from images using Brain Extraction Tool FSL-BET [43] by optimizing the fractional intensity threshold and reducing image bias and residual neck voxels. A study-specific grey matter template was then created using the FSL-VBM library and relevant protocol, found at http://fsl.fmrib.ox.ac.uk/fsl/fslwiki/FSLVBM [45]. In this step, all brain-extracted images were segmented to grey matter (GM), white matter (WM) and cerebrospinal fluid (CSF). GM images were selected and registered to the GM ICBM-152 standard template using linear affine transformation. The registered images were concatenated and averaged and were then flipped along the x-axis, and the two mirror images were then re-averaged to obtain a first-pass, study-specific affine GM template. Second, the GM images were re-registered to this affine GM template using non-linear registration, concatenated into a 4D image which was then averaged and flipped along the x-axis. Both mirror images were then averaged to create the final symmetric, study-specific “non-linear” GM template at 2×2×2 mm^3^ resolution in standard space. Following this, all concatenated and averaged 3D GM images (one 3D image per subject) were concatenated into a stack (4D image = 3D images across subjects). Additionally, the FSL-VBM protocol introduced a compensation or modulation for the contraction/enlargement due to the non-linear component of the transformation, where each voxel of each registered grey matter image was multiplied by the Jacobian of the warp field. The modulated 4D image was then smoothed by a range of Gaussian kernels, sigma = 2, 3, 4 mm (standard sigma values in the field of MRI data analysis), which approximately resulted in full width at half maximums (FWHM) of 4.6, 7 and 9.3 mm. The various spatial smoothing kernels enabled us to explore whether classification accuracy would improve by varying the spatial smoothing kernels. The MRI preprocessing module was applied to AD and NC data and produced two sets of four 4D images, which were called Structural MRI 0 – fully preprocessed without smoothing – as well as three fully preprocessed and smoothed datasets called Structural MRI 2, 3, 4, which were used in subsequent classification steps.

## 5 Results and Discussion

### 4.1 rs-fMRI Pipeline

The preprocessed rs-fMRI time series data were first loaded into memory using neuroimaging package Nibabel (http://nipy.org/nibabel/) and were then decomposed into 2D (x,y) matrices along z and time (t) axes. Next, the 2D matrices were converted to lossless PNG format using the Python OpenCV (opencv.org). The last 10 slices of each time course were removed since they included no functional information. Also, any slices with sum of pixel intensities equal to zero were ignored. During the data conversion process, a total of 793,800 images were produced, including 270,900 Alzheimer’s and 522,900 normal control PNG samples. Equation 4 describes the conversion of a given slice to a PNG sample which applies to every subject’s time course.

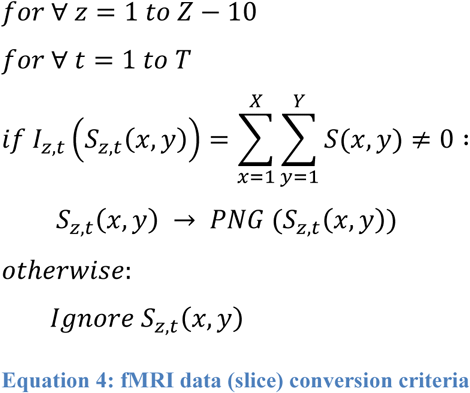

Where x, y and z are spatial dimensions (from 1 to X, Y, Z, respectively), t is a time point of a given fMRI time course with T points, S_z,t_(x, y) is a given slice with a dimension of (x, y) for the position of (z, t) and I_z_,_t_ represent the intensity function of S_z,t_(x, y). PNG represents the lossless PNG transformation function.

In the data conversion step, the 4D time courses of subjects were randomly shuffled, and five random datasets were created in order to repeat training and testing of the CNN classifier (fivefold cross-validation against all of the data). The random datasets were labeled for binary classification, and 75% of the images were assigned to the training dataset, while the remaining 25% were used for testing purposes. The training and testing images were resized to 28×28 pixels and were then converted to the Lightning Memory-Mapped Database (LMDB) for high throughput for the Caffe Deep Learning platform [12] used for this classification experiment. The adopted LeNet architecture was adjusted for 30 epochs and initialized for Stochastic Gradient Descent with gamma = 0.1, momentum = 0.9, learning rate = 0.01, weight_decay = 0.005, and the step learning rate policy dropped the learning rate in steps by a factor of gamma every stepsize iteration. The mean of images was calculated and subtracted from each image.

Training and testing of Caffe models were performed and were repeated five times on the Amazon AWS Linux G2.8×large, including four high-performance NVIDIA GPUs, each with 1,536 CUDA cores and 4GB of video memory and 32 High Frequency Intel Xeon E5-2670 (Sandy Bridge) vCPUs with 60 GB memory overall. An average accuracy rate of 99.9986% was obtained for five randomly shuffled datasets using the adopted LeNet architecture shown in Table 3. Alternatively, the first set of five randomly shuffled datasets was resized to 256×256 and was then converted to LMDB format. The adopted GoogleNet was adjusted for 30 epochs and initialized with the same parameters mentioned above, and the experiment was performed on the same GPU server. An accuracy testing rate of 100%, as reported in the Caffe log file, was achieved (in practice, Caffe rounded the accuracy up after the seventh decimal), as shown in Table 3. A very high level of accuracy of testing rs-fMRI data was obtained from both of the adopted LeNet and GoogleNet models. During the training and testing processes, the loss of training, loss of testing and accuracy of testing data were monitored. In Figure 4, the accuracy of testing and the loss of testing of the first randomly shuffled dataset are presented for the adopted LeNet and GoogleNet models, respectively. To confirm the reproducibility of the results, the entire process described above was repeated on the same server using NVIDIA DIGITS Caffe (the Deep Learning GPU Training System) and the identical results were replicated.

**Figure 4.**
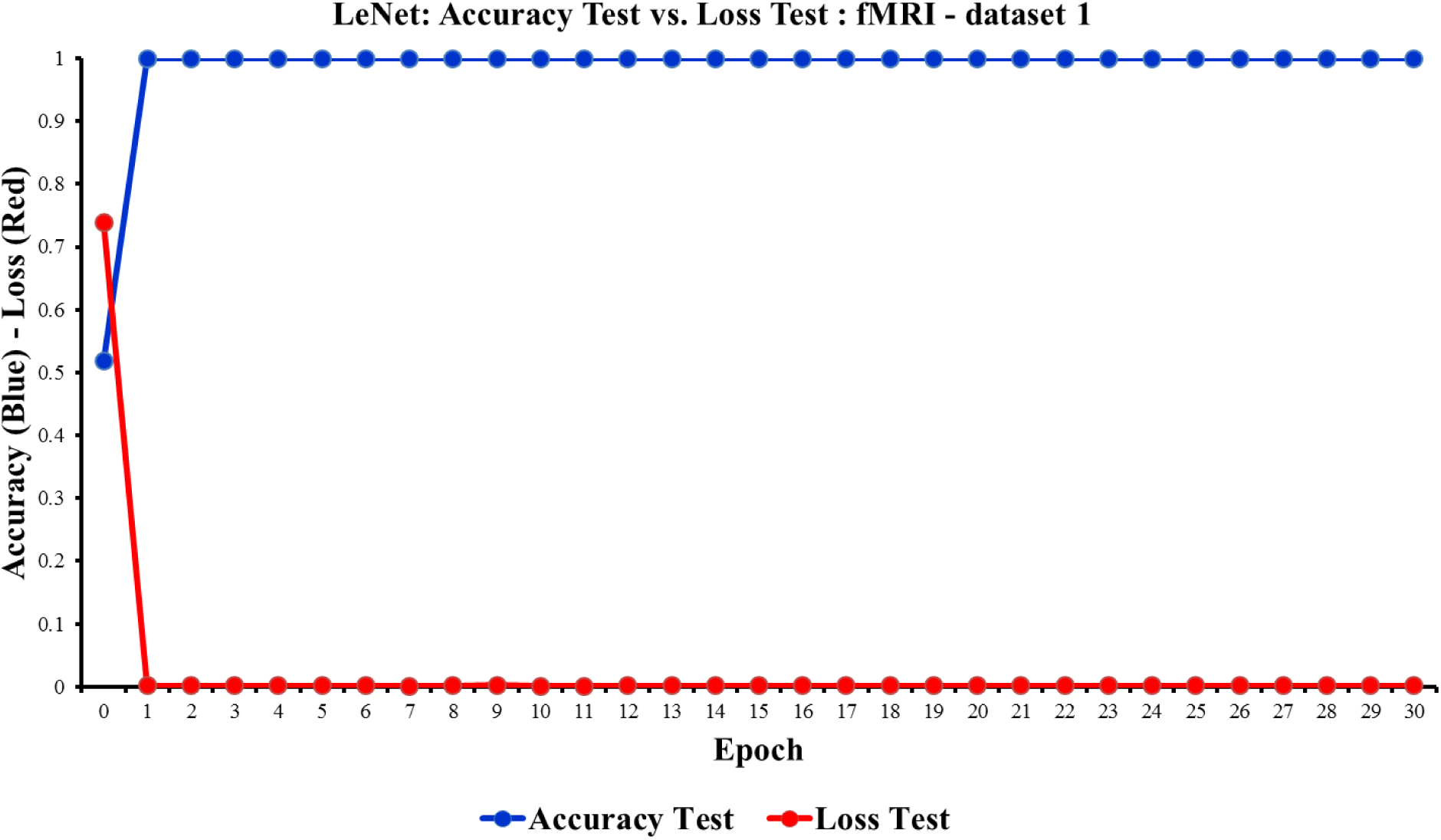
**The accuracy and loss of the first testing dataset are shown over 30 epochs. As seen, the accuracy of testing data reached almost 99.99%, and the loss of testing data dropped down to zero in the LeNet classifier.**

**Figure 5.**
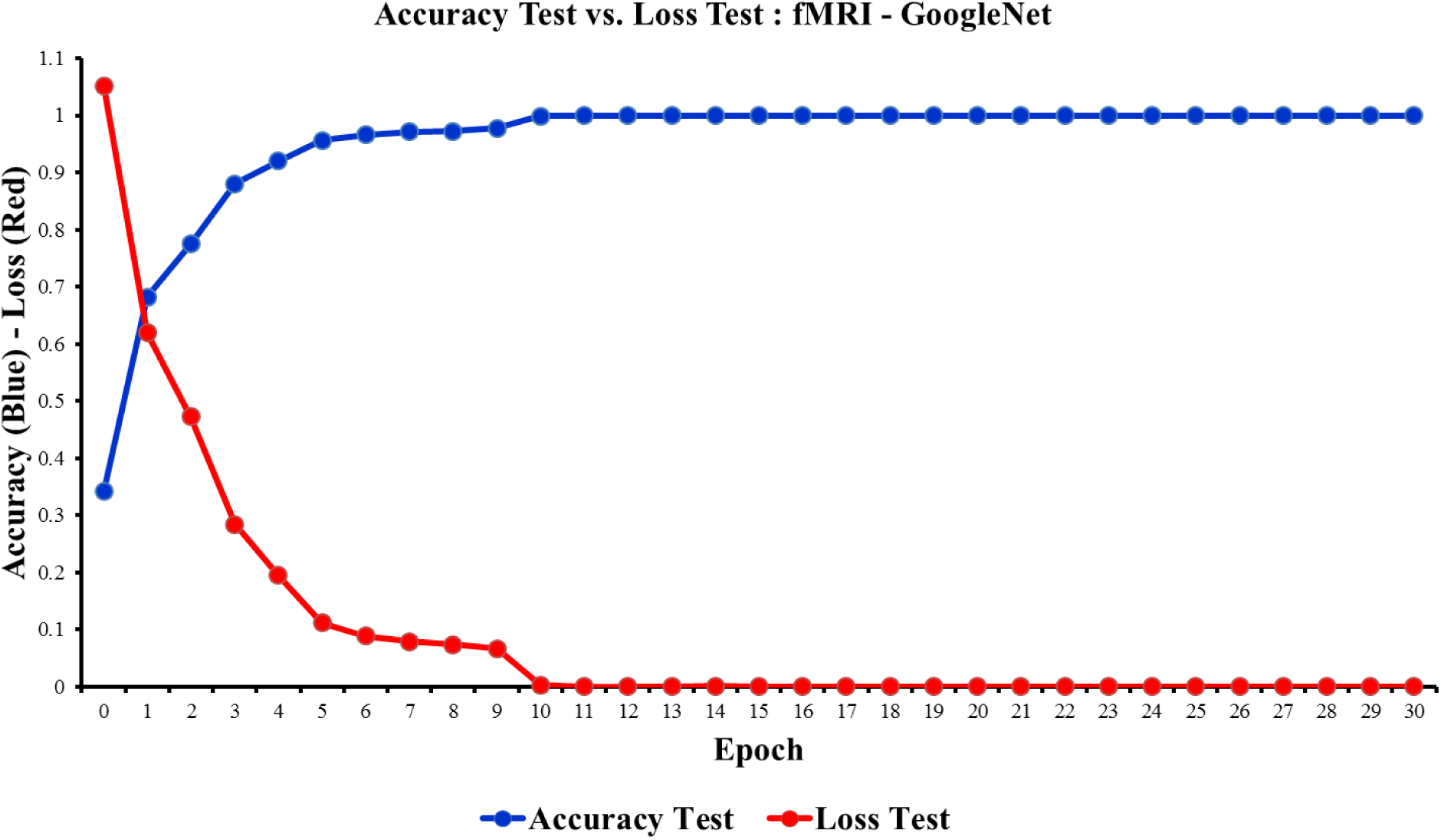
**Adopted GoogleNet training and testing resulted in a very high level of accuracy of almost 100%. As seen, the loss of testing approached zero in the 10^th^ epoch. The accuracy rates of both the LeNet and GoogleNet networks were close. However, the final accuracy of GoogleNet was slightly better than the LeNet model.**

### 5.2 Structural MRI Pipeline

The preprocessed MRI data were then loaded into memory using a similar approach to the fMRI pipeline and were converted from NII to lossless PNG format using Nibabel and OpenCV, which created two groups (AD and NC) × four preprocessed datasets (MRI 0,2,3,4). Additionally, the slices with zero mean pixels were removed from the data. The conversion criteria are similar to Equation 4 but without removing any slice from the end of 3D image and *t* represents the subject number in the stack of structural MRI data. This step produced a total number of 62,335 images, with 52,507 belonging to the AD group and the remaining 9,828 belonging to the NC group per dataset. The data were next converted to the LMDB format and resized to 28×28 pixels. The adopted LeNet model was set for 30 epochs and initiated for Stochastic Gradient Descent with gamma = 0.1, momentum = 0.9, base learning rate = 0.01, weight_decay = 0.0005, and a step learning rate policy dropping the learning rate in steps by a factor of gamma every stepsize iteration. Next, the model was trained and tested by 75% and 25% of the data for four different datasets. The training and testing processes were repeated five times on Amazon AWS Linux G2.8×large to ensure the robustness of the network and achieved accuracy. The average of accuracies was obtained for each experiment separately, as shown in Table 3. The results demonstrate that a high level of accuracy was achieved in all of the experiments, with the highest accuracy rate of 98.79% achieved for the structural MRI dataset, which was spatially smoothed by sigma = 3 mm. In the second run, the adopted GoogleNet model was selected for binary classification. In this experiment, the preprocessed datasets were converted to LMDB format and resized to 256×256. The model was adjusted for 30 epochs using Stochastic Gradient Descent with gamma = 0.1, momentum = 0.9, base learning rate = 0.01, weight_decay = 0.0005, and a step learning rate policy. The GoogleNet model resulted in a higher level of accuracy than the LeNet model, with the highest overall accuracy rate of 98.8431% achieved for MRI 3 (smoothed by sigma = 3mm). However, the accuracy rate of the unsmoothed dataset (MRI 0) reached 84.5043%, which was lower than the similar experiment with the LeNet model. This result may demonstrate the negative effect of interpolation on unsmoothed data, which may in turn strengthen the concept of spatial smoothing in MRI data analysis.

In practice, most classification questions address imbalanced data, which refers to a classification problem in which the data are not represented equally and the ratio of data may exceed 4 to 1 in binary classification. In the MR analyses performed in this study, the ratio of AD to NC images used for training the CNN classifier was around 5 to 1. To validate the accuracy of the models developed, a new set of training and testing was performed by randomly selecting and decreasing the number of AD images to 10,722 for training, while the same number of images – 9,828 – was used for the NC group. In the balanced data experiment, the adopted LeNet model was adjusted for 30 epochs using the same parameters mentioned above and was trained for four MRI datasets. In Table 3, the new results are identified with labels beginning with the B. prefix (Balanced). The highest accuracy rate obtained from the balanced data experiment only decreased around 1% (B. Structural MRI 3 = 97.81%) compared to the same datasets in the original training. This comparison demonstrates that the new results were highly correlated to the initial results, confirming that even a precipitous decrease in the data ratio from 5:1 to 1:1 had no impact on classification accuracy, which validated the robustness of the trained models in the original MRI classification.

**Table 3.**
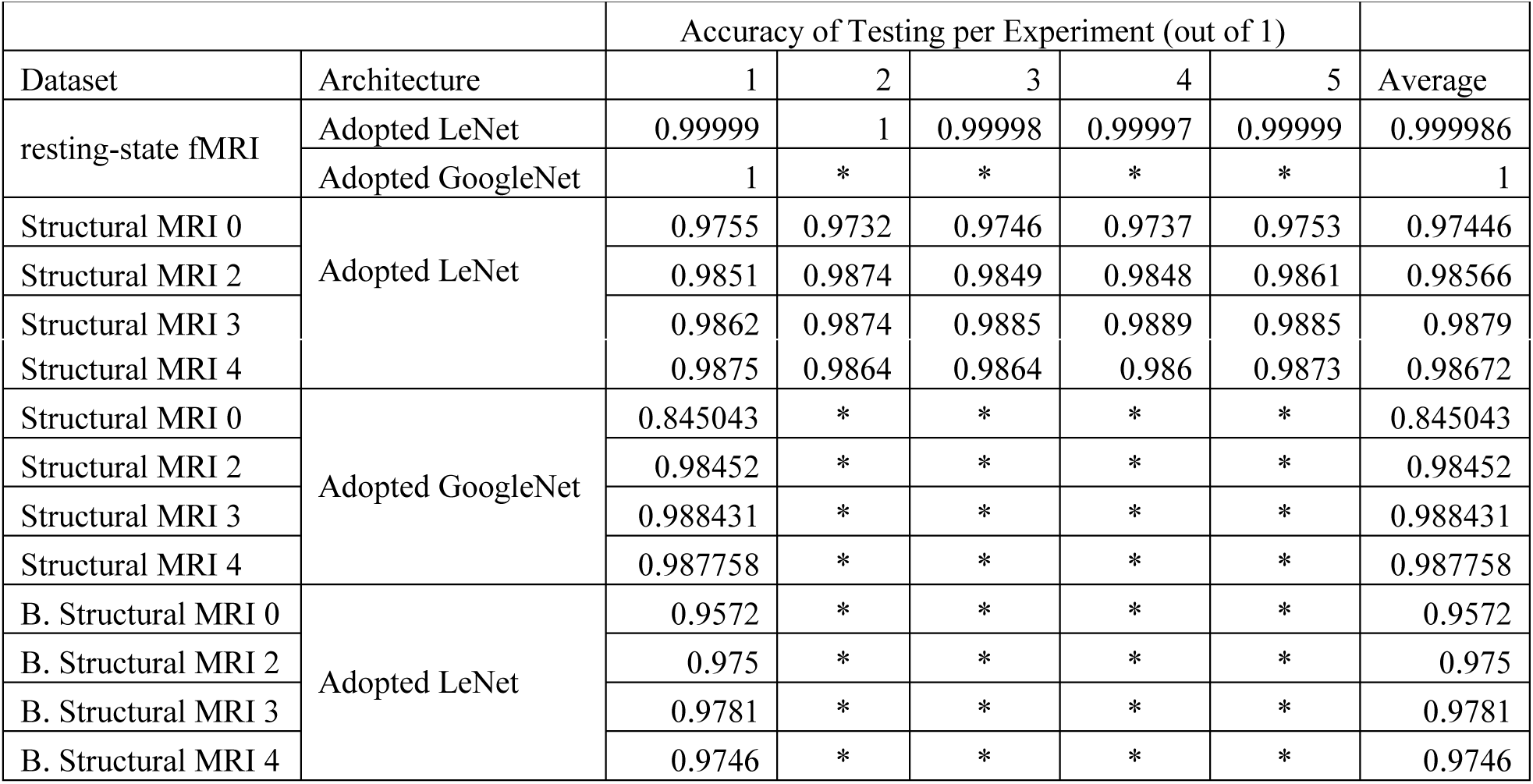
**The accuracy of testing datasets is demonstrated below. As shown, a very high level of accuracy in testing datasets was achieved in both fMRI and MRI modalities in all of the runs. The experiment of the cells with asterisks * was not required in this study. Therefore, no value was assigned. The datasets used for testing balanced data begin with the prefix B. Abbreviation: MRI 0, the structural MRI dataset without spatial smoothing. MRI 2,3,4 are the datasets spatially smoothed by Gaussian kernel sigma = 2,3 and 4 mm.**

Differentiation between subjects with Alzheimer’s disease and normal healthy control subjects (older adults) requires solid preprocessing and feature learning, which reveal functional and structural dissimilarities between Alzheimer’s damage and routine effects of age on the brain. In this study, two robust pipelines were designed and implemented that were capable of producing consistent and reproducible results. In the first block of the pipelines, extensive data preprocessing was performed against fMRI and MRI data, which removed potential noise and artefacts from the data. Next, a convolutional layer of CNN architecture consisting of a set of learnable filters, and which also serves as a shift and scale invariant operator, extracted low- to mid-level features (as well as high-level features in GoogleNet). In the fMRI pipeline, both adopted LeNet and GoogleNet architecture were trained and tested by a massive number of images created from 4D fMRI time series. Furthermore, removal of non functional brain images from data improved the accuracy of recognition when compared to previous experience [46]. In the MRI pipeline, four sets of images (smoothed with different kernels) were used to train and test the CNN classifier to ensure that the best preprocessed data were employed to achieve the most accurate trained model. The results demonstrate that spatial smoothing with an optimal kernel size improves classification accuracy (Figure 6). Certain differences in image intensity (Figure 7), brain size of AD and NC subjects, and lack of signals in brain regions of AD samples, such as the frontal lobe, are strong evidence in support of the success of the pipelines.

**Figure 6.**
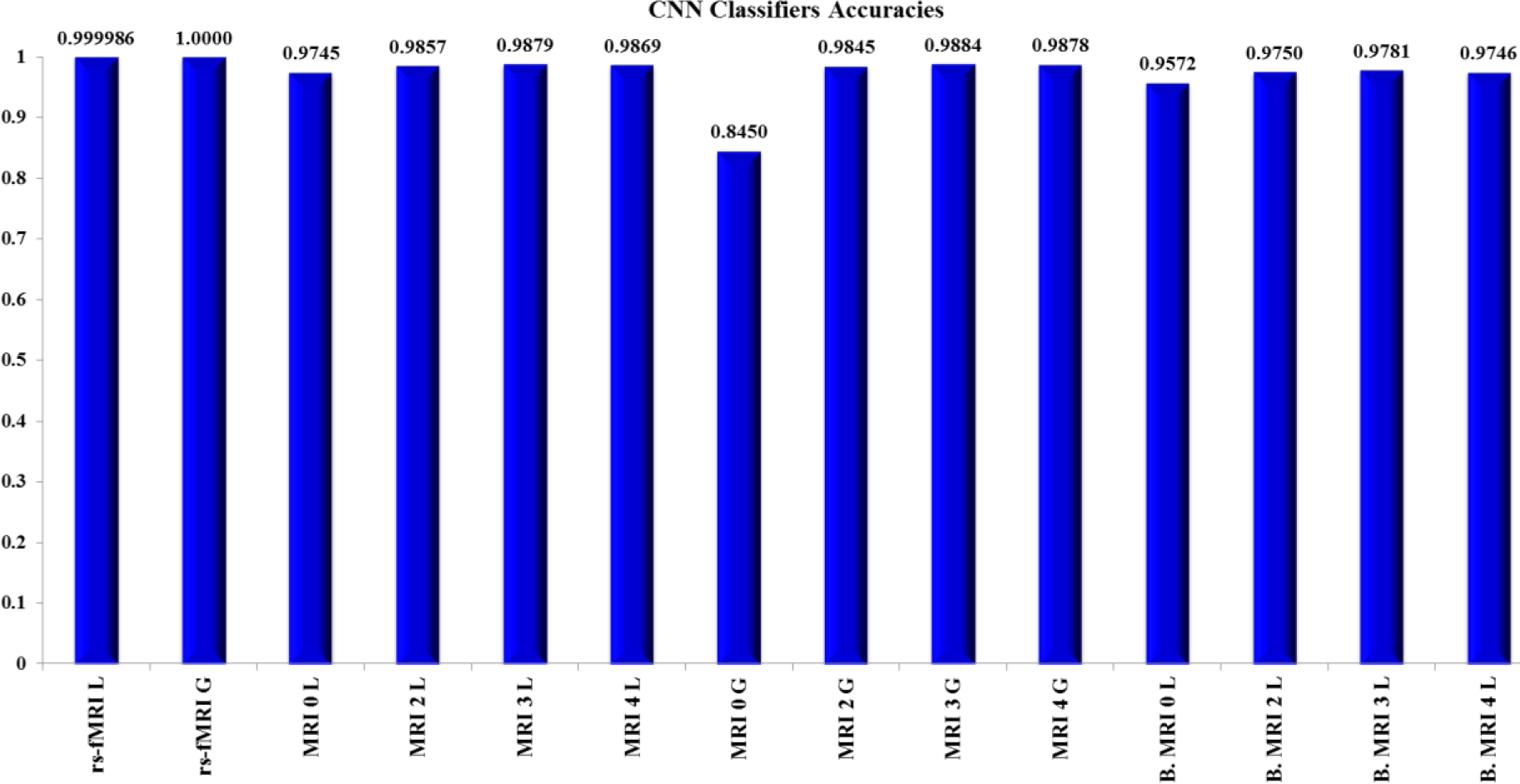
**A total of 14 values, including five averaged accuracies and nine single accuracies from a total of 29 training CNN-based classifiers (adopted LeNet and GoogleNet), are demonstrated. Almost perfect accuracy was achieved using the ADNI fMRI data from both models. Additionally, the ADNI MRI data were successfully classified with an accuracy rate approaching 99%. These results demonstrate that CNN-based classifiers are highly capable of distinguishing between AD and NC samples by creating low- to high-level shift and scale invariant features. The results also demonstrate that in MRI classification, spatially smoothed data with sigma = 3 mm produced the highest accuracy rates.**

**Figure 7.**
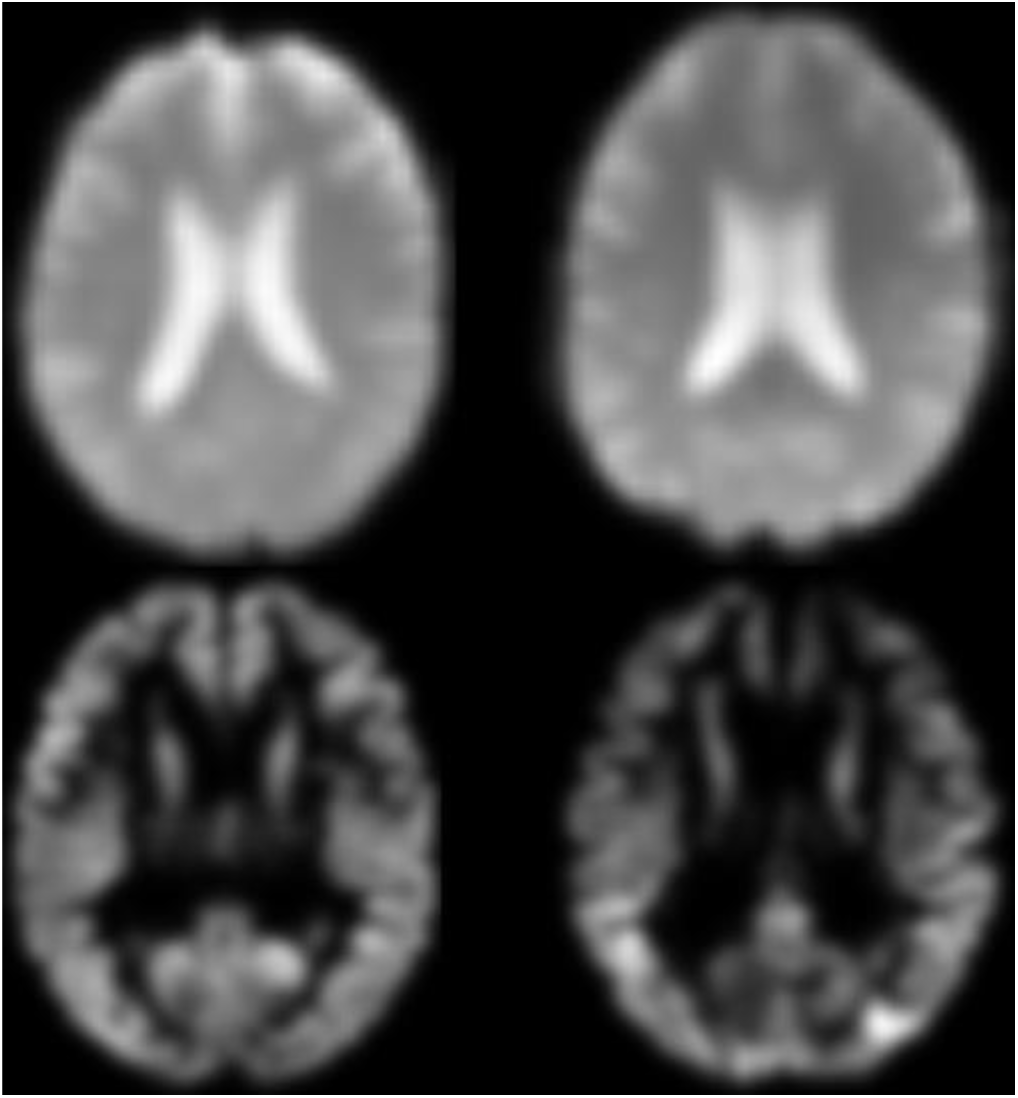
**A middle cross-section of fMRI data (22, 27, 22) with thinness of 4 mm, representing a normal healthy brain (top-left), and an Alzheimer’s brain are shown (top-right). A middle cross-section of structural MRI (45, 55, 45) with thickness of 2 mm, representing a normal brain (bottom-left), and an Alzheimer’s subject (bottom-right) are also demonstrated. In both fMRI and MRI modalities, different brain patterns and signal intensities are identified.**

A common strategy employed is to visualize the weights of filters to interpret the conv layer results. These are usually most interpretable on the first conv layer, which directly examines the raw pixel data, but it is also possible to find the filter weights deeper in the network. In a well-trained network, smooth filters without noisy patterns are usually discovered. A smooth pattern without noise is an indicator that the training process is sufficiently long, and likely no overfitting occurred. In addition, visualization of the activation of the network’s features is a helpful technique to explore training progress. In deeper layers, the features become more sparse and localized, and visualization helps to explore any potential dead filters (all zero features for many inputs). Filters and features of the first layer for a given fMRI and MRI trained LeNet model were visualized using an Alzheimer’s brain and a normal control brain. Figure 8 and Figure 9 demonstrate 20 filters of 5×5 pixels for fMRI and MRI models, respectively. Additionally, 20 features of 24×24 pixels in Figure 10 and Figure 11 reveal various regions of the brain that were activated in AD and NC samples.

**Figure 8.**
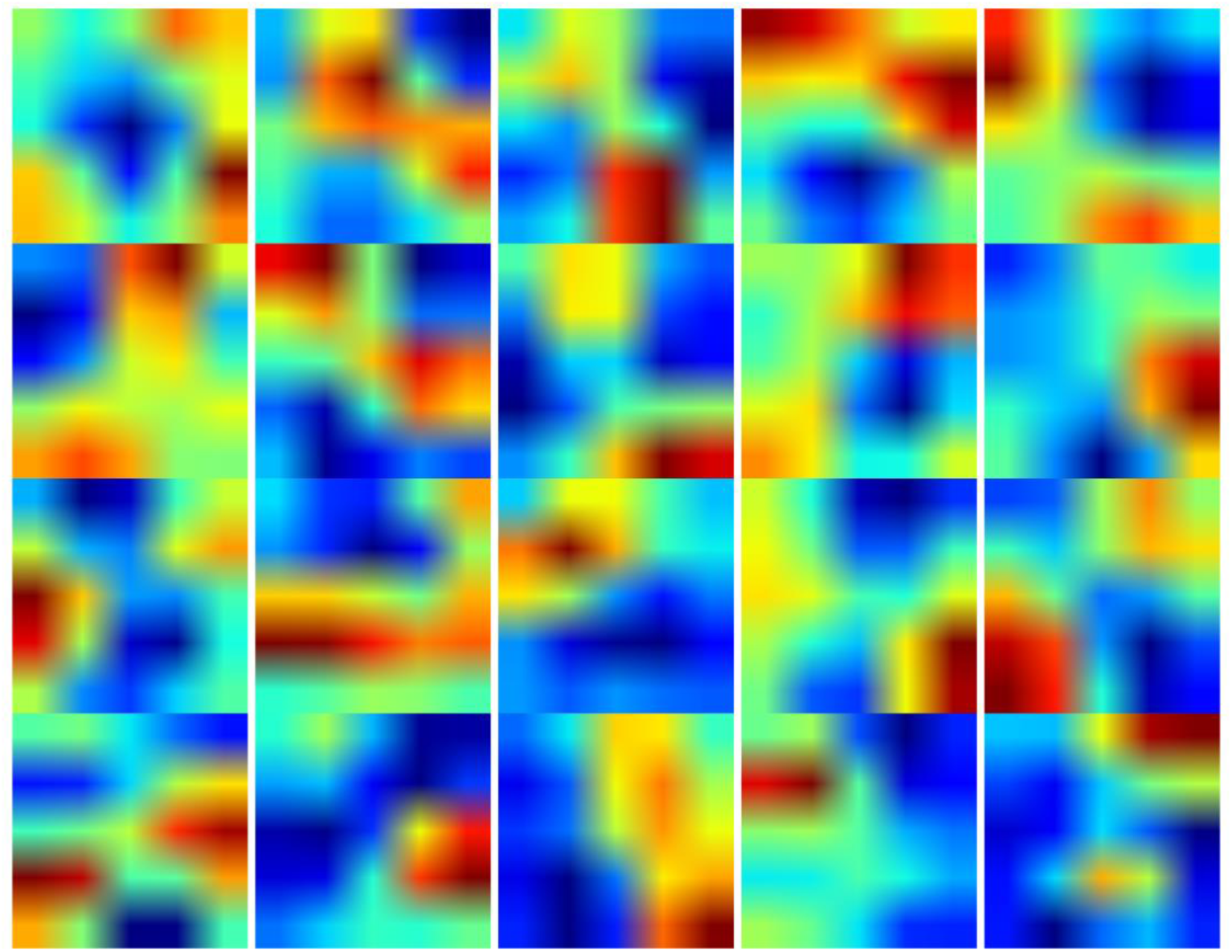
**In the first layer of LeNet in a given trained fMRI model, 20 filters of 5×5 pixels were visualized. The weights shown were applied to the input data and produced activation, or features, of a given sample.**

**Figure 9.**
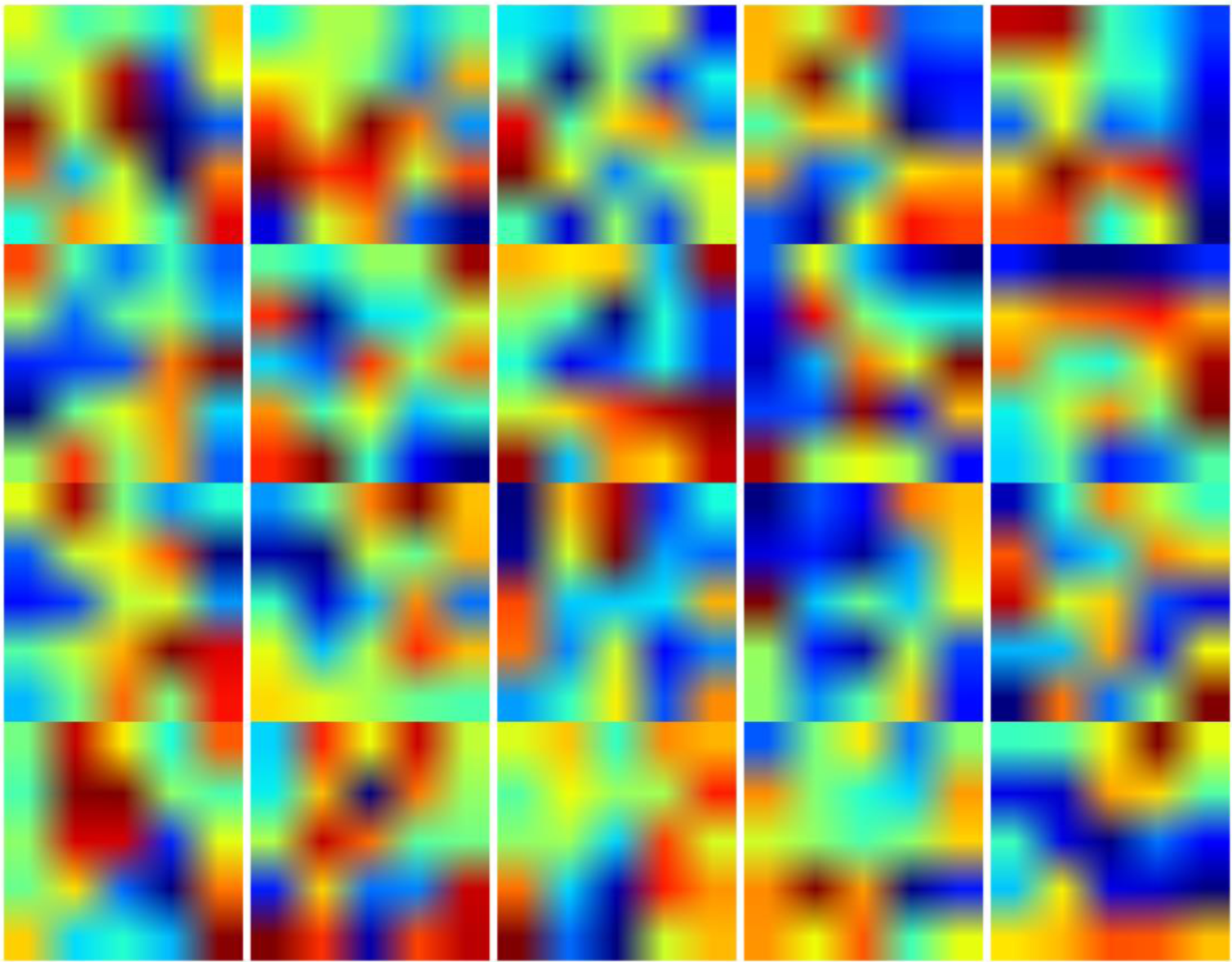
**In a trained LeNet model, 20 filters with a kernel of 5×5 were visualized for the first layer. The filters shown were generated from a model in which MRI data smoothed by sigma = 3 mm were used for training.**

**Figure 10.**
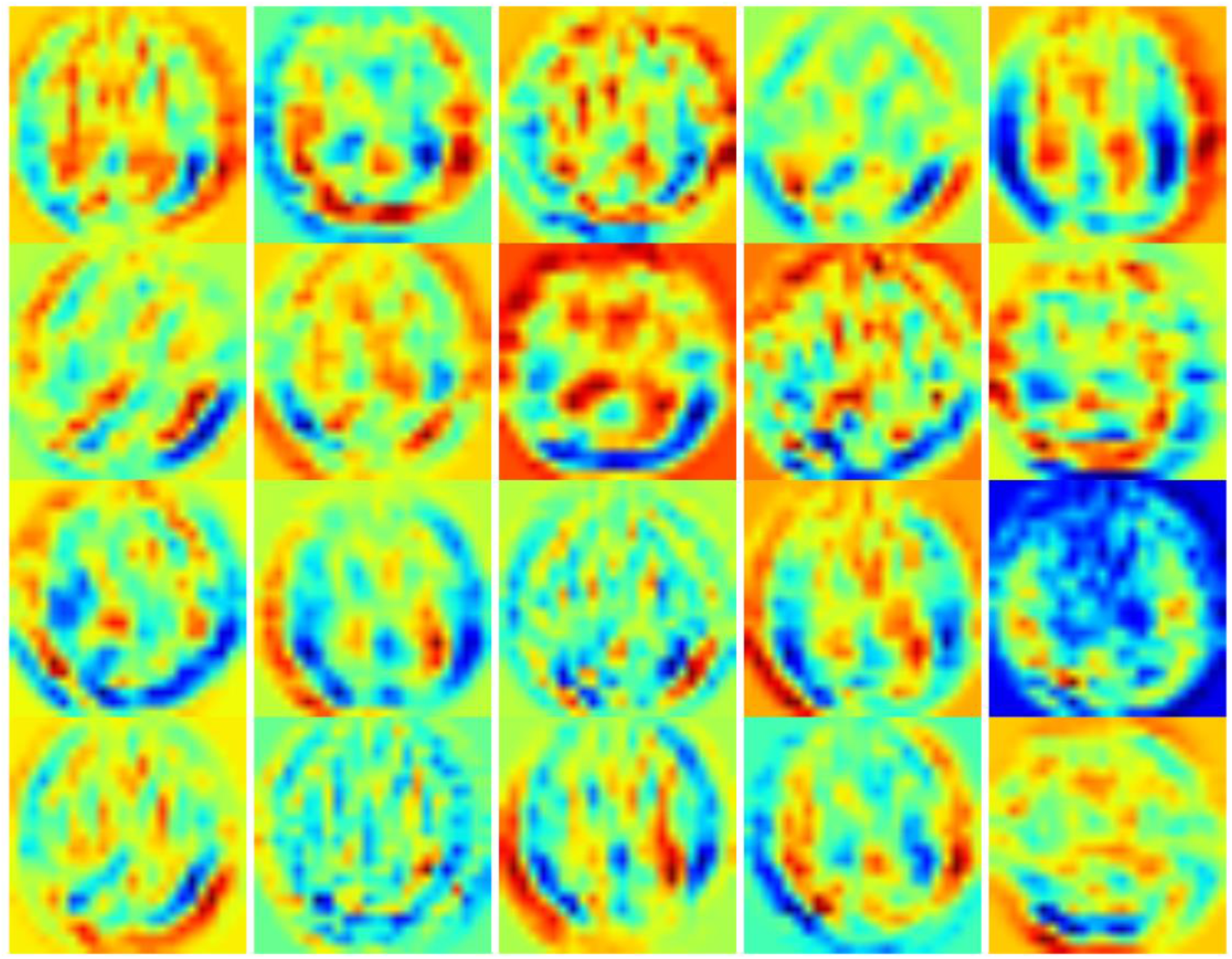
**20 activations (features) of the first layer of LeNet trained using MRI data were displayed for a given AD MRI sample (45, 55, 45). A smooth pattern without noise reveals that the model was successfully trained.**

**Figure 11.**
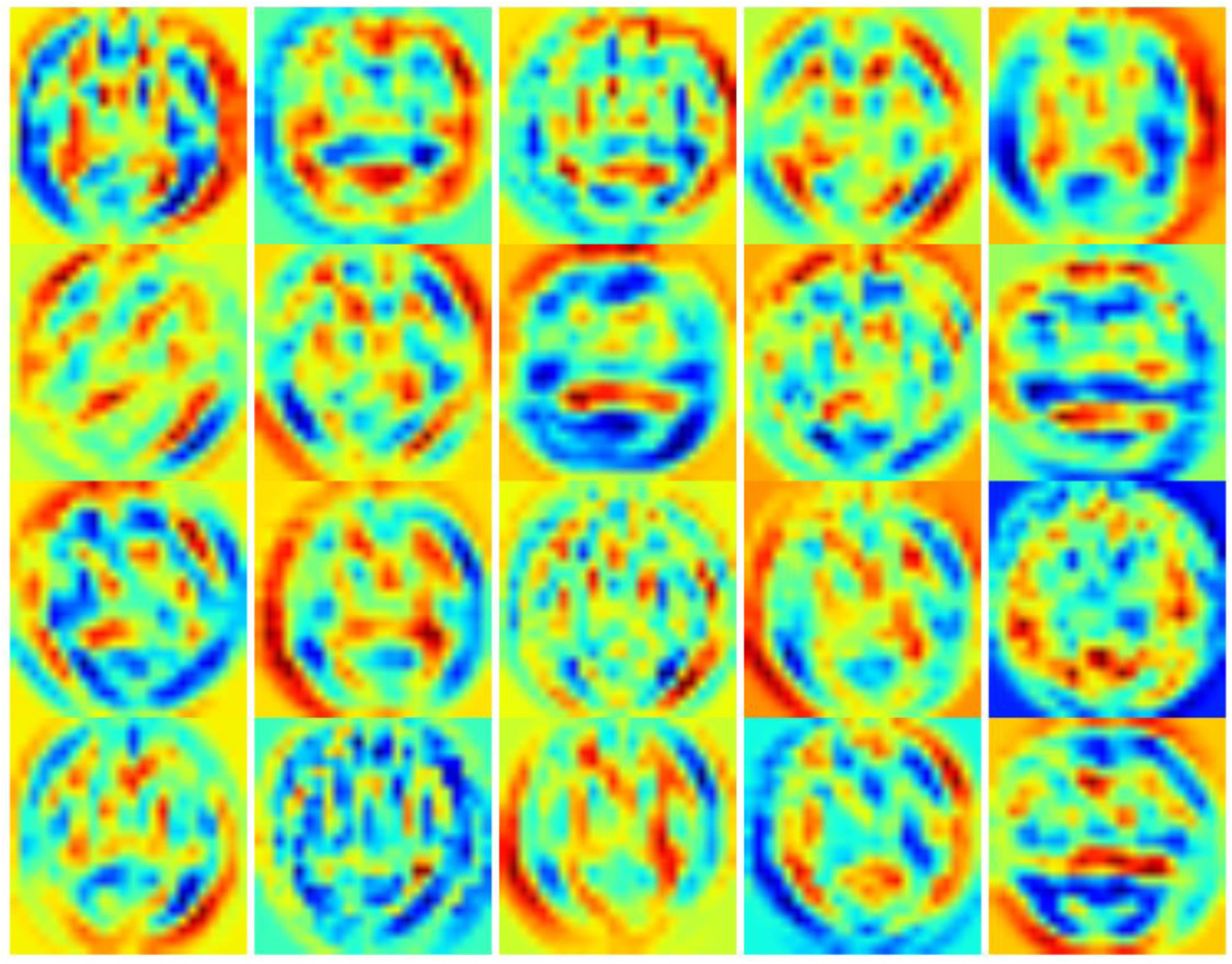
**Features of the first layer of the same MRI trained model were displayed for a normal control (NC) brain slice (45, 55, 45). A basic visual comparison reveals significant differences between AD and NC samples.**

## 6 Subject-level recognition

To expand the application for distinguishing Alzheimer’s subjects from normal healthy brains to the clinical level, subject-level classification of AD and NC was investigated. In the clinical application, neuroscientists and neuropsychologists tend to validate a trained machine learning-based model, which is capable of classifying a given subject’s imaging data rather than slice-level recognition, which is in the interest of machine learning specialists. In order to perform subject-level classification, the rs-fMRI and four categories of structural MRI preprocessed data were considered. In this experiment, the training and testing subject-level samples were randomly selected by shuffling subjects five times, as shown in Table 1. Next, 75% and 25% of subjects were assigned to the training and testing datasets, respectively. As mentioned previously, none of the training and testing samples came from the subjects.

Then, the adopted LeNet model and GoogleNet were adjusted for 30 epochs using Stochastic Gradient Descent with gamma = 0.1, momentum = 0.9, base learning rate = 0.01, weight decay = 0.0005, and a step learning rate policy. Table 4 demonstrates the results for the five categories and five cross-validations, including one rs-fMRI and four structural MRIs. Although the accuracy rates decreased slightly in certain cases as expected, those rates were still very high and better than any accuracy rates reported in the literature, which demonstrates the robustness, reproducibility and reliability of the DeepAD pipeline. Unlike the slice-level experiment, the highest accuracy rates belonged this time to the structural MRI samples smoothed by a Gaussian kernel sigma of 3 mm (and sigma = 4) for both the adopted LeNet and GoogleNet, as shown in Figure 12. This strict validation process also revealed that the implemented pipeline was capable of extracting the pattern and distinguishing AD from NC brains among uniformly correlated or semi-correlated samples that had been preprocessed and normalized. On the other hand, as the variety of samples used in the training was less than in the previous method, a slight drop in accuracy rate was expected. In real-world applications of machine learning, the validation process should be as strict as possible, but training the classifier by using a wide range of data is a must. Otherwise, the trained model will be unable to make generalizations.

**Table 4.**
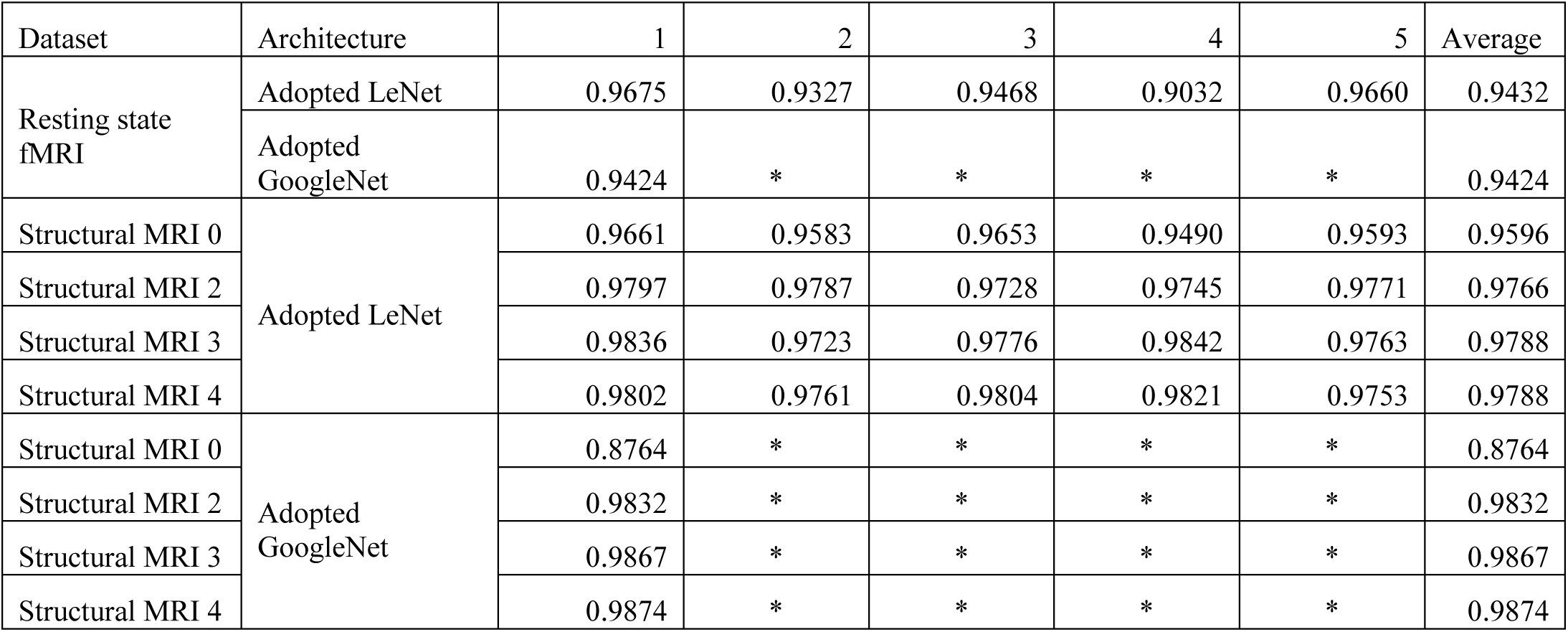
**Subject-level accuracy rates of validation results for adopted LeNet and GoogleNet against rs-fMRI and structural MRI samples**

**Figure 12.**
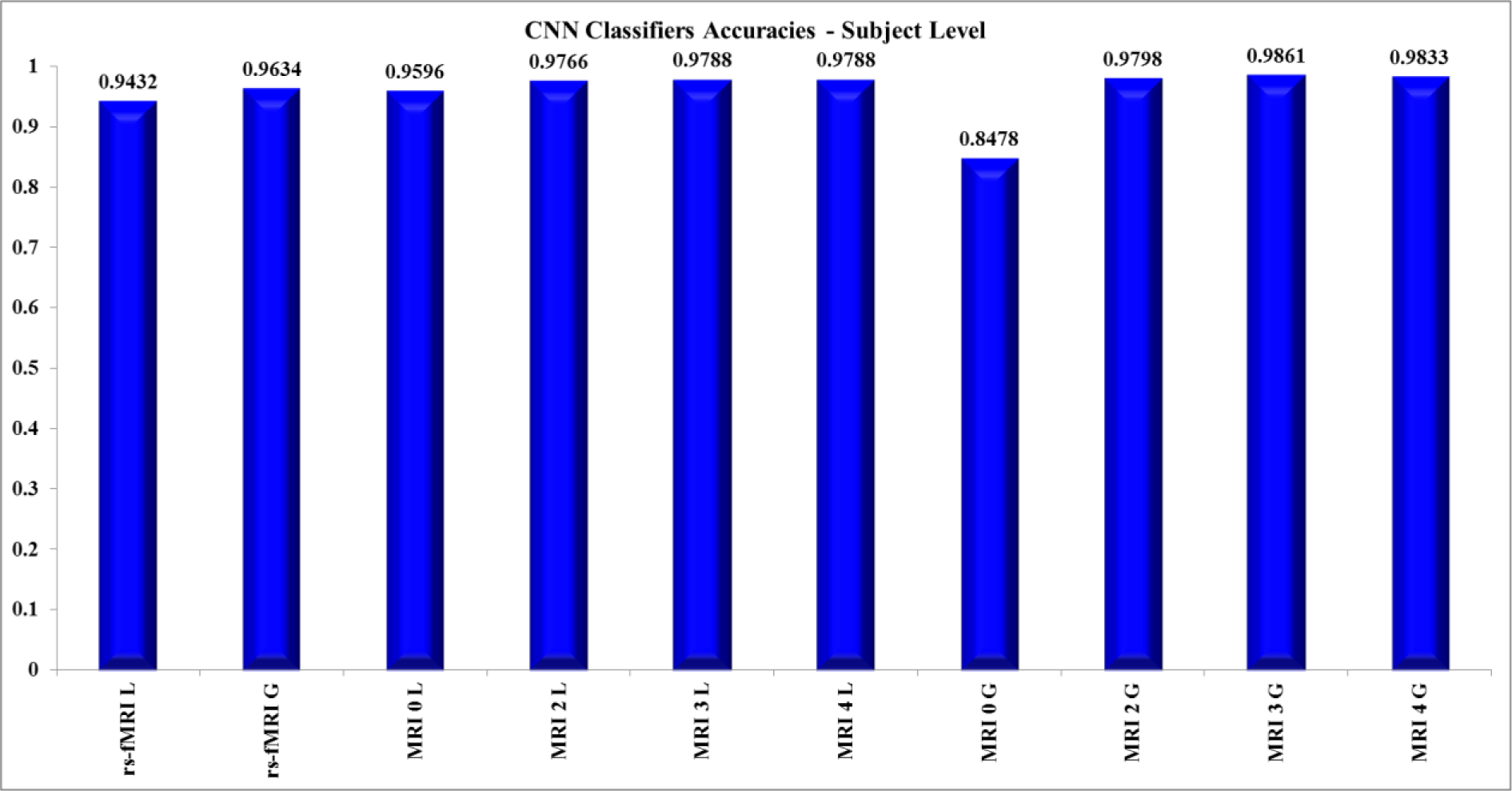
**The averaged accuracy rates of validation (Testing) samples for 10 categories of rs-fMRI and structural MRI and adopted LeNet and GoogleNet. As seen in the figure, all categories resulted in very high accuracy rates except for the structural MRI dataset without smoothing.**

### 6.1 ROC Curve

The receiver operating characteristic (ROC) curves were illustrated to validate the performance of the binary classifiers trained using the DeepAD pipeline. For each trained LeNet model in both modalities, the true positive, true negative, false positive and false negative rates were calculated for 10 thresholds from 0 to 1 with steps of 0.1 (0:1:0.01). Next, for each classifier, the sensitivity and specificity were calculated using Equation 5.

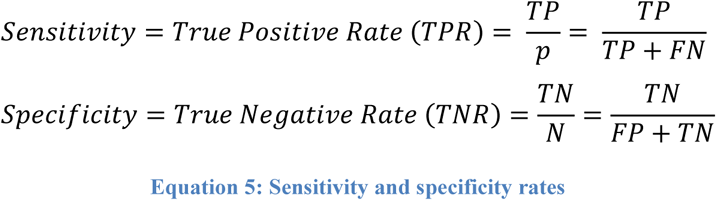

Next, for each category of classification, the TPR and TNR rates were averaged for visualization purposes. As shown in Figure 13 and Figure 14, the performance of classifiers inclined toward the ideal classification and diagnostic performance, and was beyond the random guess.

**Figure 13.**
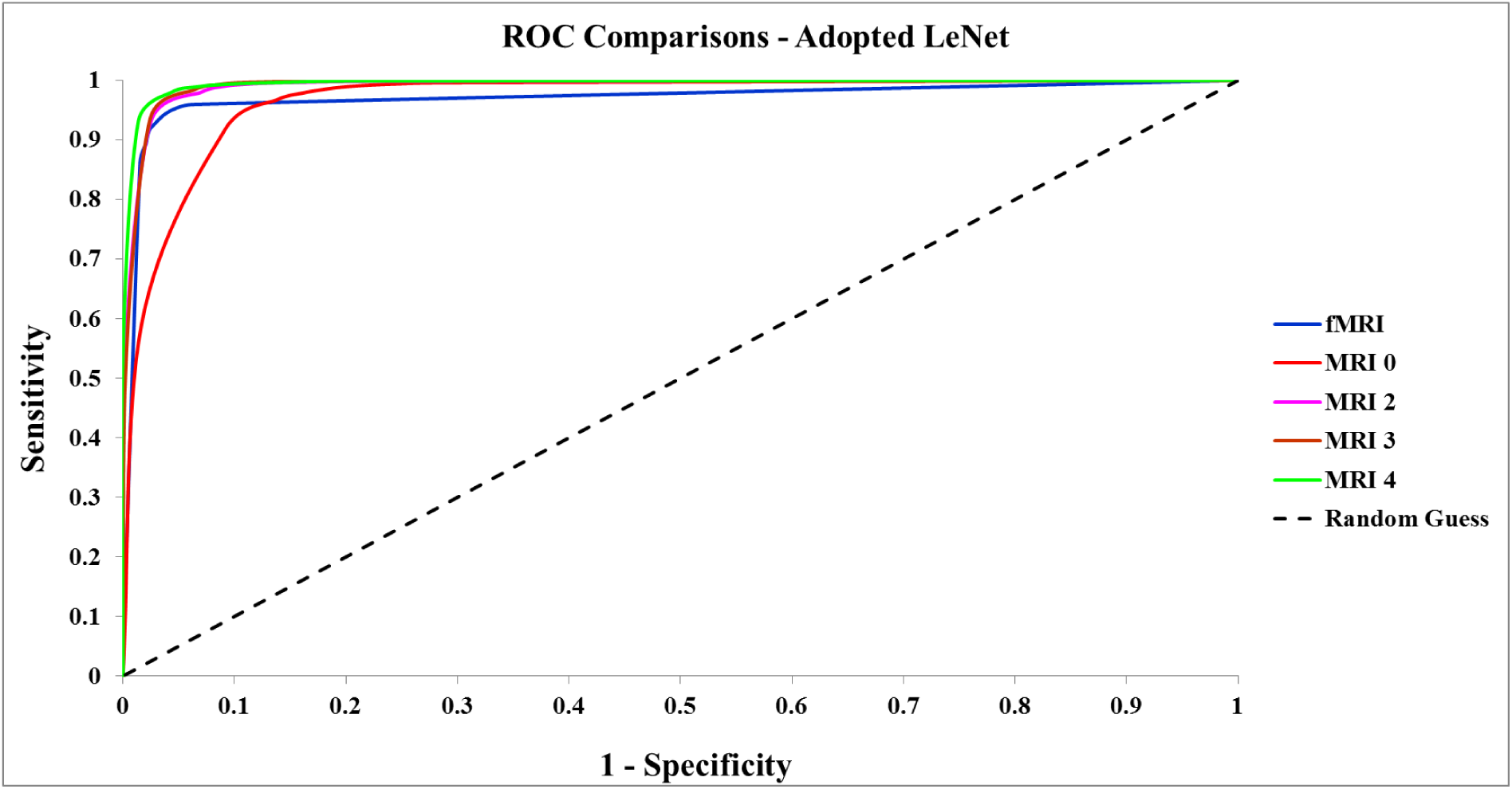
**To calculate ROC curves, the averaged sensitivity and 1-specificity were used for five categories of classification which included the adopted LeNet models of rs-fMRI and structural MRI smoothed by four different Gaussian kernels. Comparison of ROC curves indicated that the performance of 1) each classifier and 2) DeepAD pipeline were beyond random guess and tended towards the perfect binary classification.**

**Figure 14.**
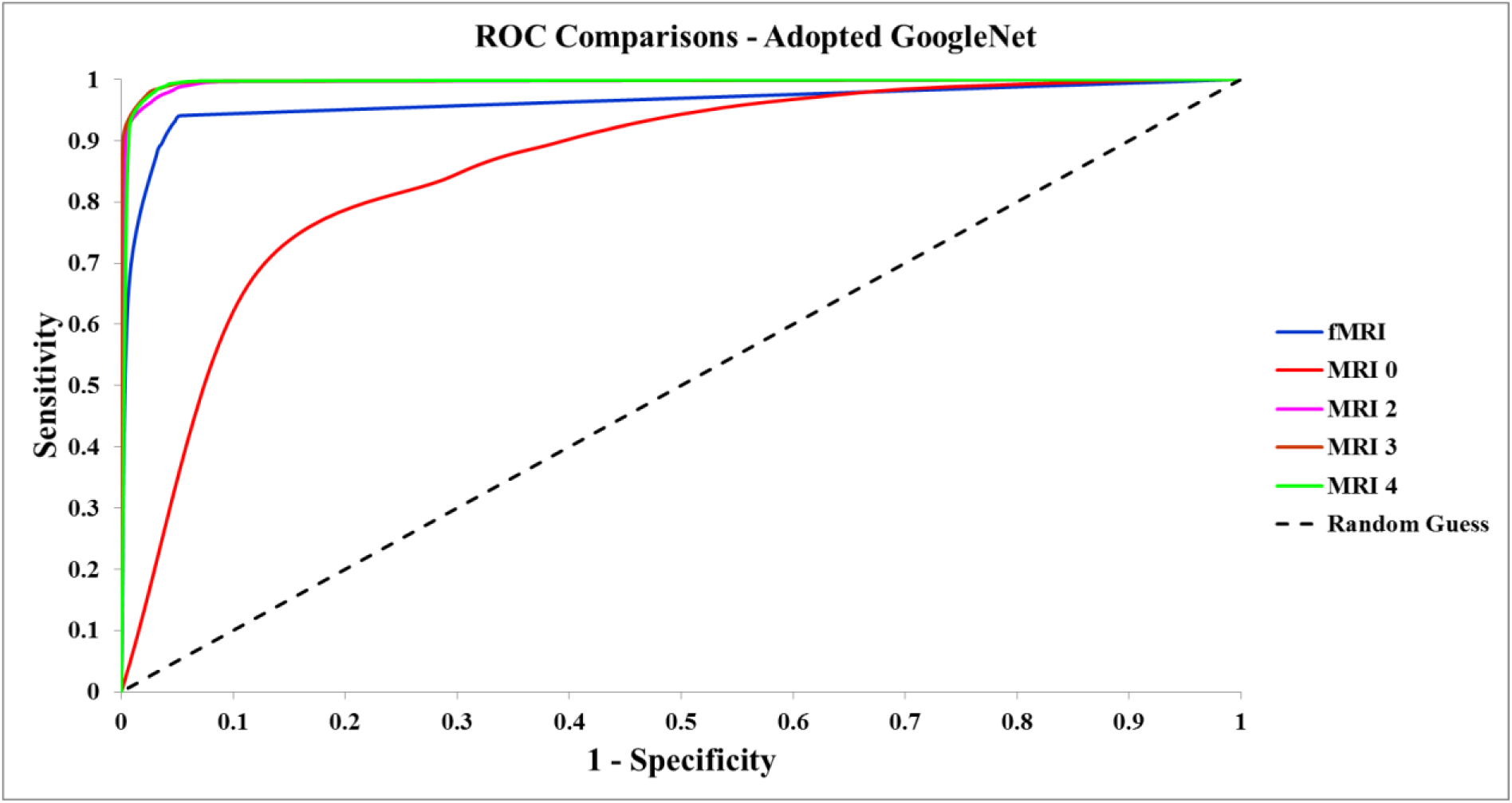
**ROC curves drawn for Sensitivity over 1-Specificity illustrate the performance of the adopted GoogleNet classifiers. The MRI dataset smoothed by Gaussian filters of 3 and 4 mm show the best performance. The MRI 2 and fMRI also indicate that the classification was very well performed; however, the MRI dataset without spatial smoothing could not compete with other classifiers.**

## 7 Decision making: Majority rule

A decision-maker algorithm is a means for discovering the best choice among a list of alternatives based on the preferred single criterion or multi-criteria and values. In a majority rule-based system, a binary decision is made by voting for the class or group that has the highest number of candidates. A decision-making algorithm was developed to classify and assign a given subject to the Alzheimer’s or normal control group by voting for the majority based on the assumption that a given subject with more AD slices is potentially AD while one with more NC slices is probably NC. In each subject, the number of slices for each class was calculated. Then, the number of slices for each class within a subject was compared, and the class with most slices was presented as the candidate. In the case of two classes having identical probability, which did not occur in the actual experiment, the decision maker would recommend a decision based on the clinical tests and measures. This algorithm enabled the entire pipeline, especially the subject-level pipeline, to produce a highly accurate prediction while also compensating for the slight loss during training and validation. The details of the decision making algorithm are as follows:

Step 1: Calculate the number of AD or NC slices in a given class where i is slice number.

~~~
        For all i = 1, … do
                If slice (i) ∈ AD
                        Counter of AD += 1
                Else if slice (i) ∈ NC
                        Counter of NC += 1
~~~

Step 2: Calculate the probability of each class by dividing the number of slices in a class by the total number of slices.

~~~
        Probability AD = Counter of AD / Total number of slices in the subject
        Probability NC = Counter of NC / Total number of slices in the subject
~~~

Step 3: Compare the two probabilities for the AD and NC classes

~~~
        If Probability AD = Probability NC
                Flag = Decision is to be made based on clinical measures
        If Probability AD > Probability NC
                Flag = AD
        Else
                Flag = NC
~~~

Step 4: Vote for the majority and assign the label of ‘majority group’ to the given subject.

~~~
        Assign Flag to the given subject
~~~

The decision-maker algorithm was applied to all adopted LeNet models for both rs-fMRI and structural MRI. The final results shown in Table 5 indicate a significant improvement in the accuracy rate of subject-level recognition. Most of the subject-level accuracies reached a rate of 100% after the decision making algorithm was applied. In other words, the post-classification method fully stabilized the very well trained deep learning mode by correcting a very small number of slices. Figure 15 showed the effect of the decision-making algorithm on subject-level LeNet trained models.

**Table 5.** **Accuracy Rate after decision making algorithm applied to subject-level LeNet models.**

| Model/Exp | LeNet 1 | LeNet 2 | LeNet 3 | LeNet 4 | LeNet 5 | GoogleNet 1 |
| --- | --- | --- | --- | --- | --- | --- |
| fMRI | 1 | 1 | 0.9706 | 0.92105263 | 1 | 0.9744 |
| MRI 0 | 1 | 0.9709 | 0.9942 | 0.9651 | 0.9826 | 0.9186 |
| MRI 2 | 1 | 1 | 1 | 1 | 1 | 1 |
| MRI 3 | 1 | 1 | 1 | 1 | 1 | 1 |
| MRI 4 | 1 | 1 | 1 | 1 | 1 | 1 |

**Figure 15.**
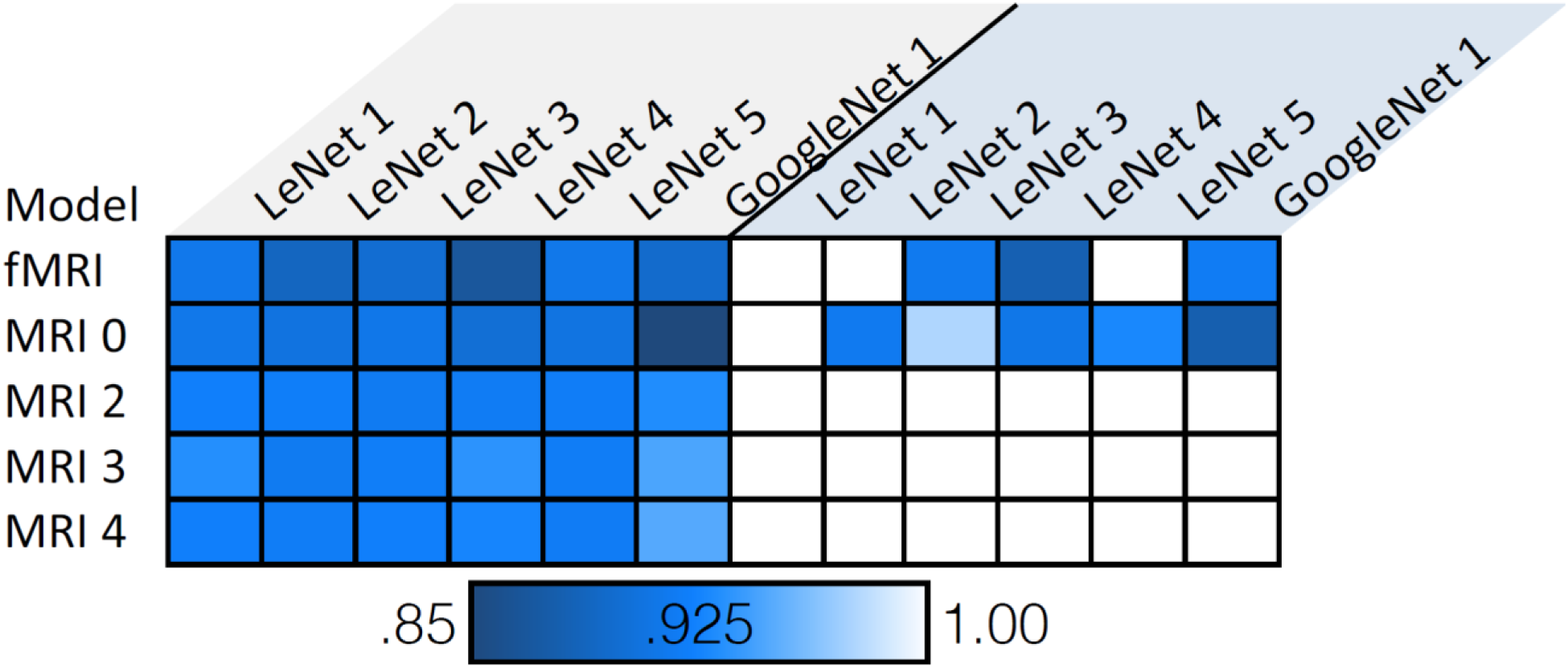
****The accuracy of five categories of adopted LeNet-based models for five repetitions as well as the adopted GoogleNet-trained networks were extracted from Table 3 and visualized in the figure on the left. After applying the decision-making algorithm, the accuracy for each experience was calculated and the results were extracted from Table 4 and visualized (right-hand figure). As shown above, the decision maker significantly improved the performance and resulted in perfect classification in most cases of the subject-level validation experiment.****

In future works, however, it may be feasible to develop a statistical model to score a given subject by defining the probabilities extent to which each subject belongs to each class based on the current trained model.

## 8 Conclusion

We presented two robust pipelines using extensive preprocessing modules and deep learning-based classifiers of MRI and fMRI data to distinguish brains affected by Alzheimer’s disease from normal healthy brains in older adults. Scale- and shift-invariant low- to high-level features were extracted from a massive volume of whole brain data using convolutional neural network architecture, resulting in a highly accurate and reproducible predictive model. In this study, the achieved accuracy rates for both MRI and fMRI modalities, as well as LeNet and GoogleNet state-of-the-art architecture, proved superior to all previous methods employed methods in the literature. Furthermore, fMRI data were used to train a deep learning-based pipeline for the first time. This successful and cutting-edge deep learning-based framework has implications for numerous applications in classifying brain disorders in both clinical trials and large-scale research studies. This study also demonstrated that the developed pipelines served as fruitful algorithms in characterizing multimodal MRI biomarkers. Lastly, the subject-level classification designed for clinical purposes enabled us to assess the robustness of the fMRI and MRI pipelines followed by a decision making algorithm, verifying that they both almost perfectly distinguish Alzheimer’s patients from healthy normal brains. In conclusion, the proposed methods demonstrated the potential for predicting stages in the progression of Alzheimer’s disease and classifying the effects of aging in the normal brain, which future studies may address.

## 9 Acknowledgment

Data collection and sharing for this project was funded by the Alzheimer's Disease Neuroimaging Initiative (ADNI) (National Institutes of Health Grant U01 AG024904) and DOD ADNI (Department of Defense award number W81XWH-12-2-0012). ADNI is funded by the National Institute on Aging, the National Institute of Biomedical Imaging and Bioengineering, and through generous contributions from the following: AbbVie, Alzheimer's Association; Alzheimer's Drug Discovery Foundation; Araclon Biotech; BioClinica, Inc.; Biogen; Bristol-Myers Squibb Company; CereSpir, Inc.; Cogstate; Eisai Inc.; Elan Pharmaceuticals, Inc.; Eli Lilly and Company; EuroImmun; F. Ho mann-La Roche Ltd and it’s a liated company Genentech, Inc.; Fujirebio; GE Healthcare; IXICO Ltd.; Janssen Alzheimer Immunotherapy Research & Development, LLC.; Johnson & Johnson Pharmaceutical Research & Development LLC.; Lumosity; Lundbeck; Merck & Co., Inc.; Meso Scale Diagnostics, LLC.; NeuroRx Research; Neurotrack Technologies; Novartis Pharmaceuticals Corporation; Pfizer Inc.; Piramal Imaging; Servier; Takeda Pharmaceutical Company; and Transition Therapeutics. The Canadian Institutes of Health Research is providing funds to support ADNI clinical sites in Canada. Private sector contributions are facilitated by the Foundation for the National Institutes of Health (www.fnih.org). The grantee organization is the Northern California Institute for Research and Education, and the study is coordinated by the Alzheimer's Therapeutic Research Institute at the University of Southern California. ADNI data are disseminated by the Laboratory for Neuro Imaging at the University of Southern California.

## References

[1] Prashanthi Vemuri, David T. Jones, and Clifford R. Jack Jr., “Resting state functional MRI in Alzheimer's Disease,” Alzheimer’s research & therapy, vol. 4, pp. 1–9, 2012.

[2] Yong He, Liang Wang, Yufeng Zang, Lixia Tian, Xinqing Zhang, Kuncheng Li, and Tianzi Jiang, “Regional coherence changes in the early stages of Alzheimer’s disease: a combined structural and resting-state functional MRI study,” Neuroimage, vol. 35, no. 2, 2007.

[3] Saman Sarraf, Jian Sun, “Functional Brain Imaging: A Comprehensive Survey,” arXiv preprint arXiv:1602.02225, 2016.

[4] Cheryl Grady, Saman Sarraf, Cristina Saverino, and Karen Campbell, “Age differences in the functional interactions among the default, frontoparietal control, and dorsal attention networks.,” Neurobiology of aging, pp. 159–172, 2016.

[5] Cristina Saverino, Zainab Fatima, Saman Sarraf, Anita Oder, Stephen C. Strother, and Cheryl L. Grady, “The Associative Memory Deficit in Aging Is Related to Reduced Selectivity of Brain Activity during Encoding.,” Journal of cognitive neuroscience, 2016.

[6] M. A. Warsi, “The Fractal Nature and Functional Connectivity of Brain Function as Measured by BOLD MRI in Alzheimer’s Disease,” macsphere.mcmaster.ca, Hamilton, 2012.

[7] Cheryl L. Grady, Anthony R. McIntosh, Sania Beig, Michelle L. Keightley, Hana Burian, and Sandra E. Black, “Evidence from functional neuroimaging of a compensatory prefrontal network in Alzheimer’s disease,” The Journal of Neuroscience, vol. 23, no. 3, pp. 986–993, 2003.

[8] Cheryl L. Grady, Maura L. Furey, Pietro Pietrini, Barry Horwitz, and Stanley I. Rapoport, “Altered brain functional connectivity and impaired short-term memory in Alzheimer’s disease,” Brain, vol. 124, no. 4, pp. 739–756, 2001.

[9] Tripoliti, Evanthia E., Dimitrios I. Fotiadis, and Maria Argyropoulou, “A supervised method to assist the diagnosis and classification of the status of Alzheimer’s disease using data from an fMRI experiment,” in In Engineering in Medicine and Biology Society, 2008. EMBS 2008. 30th Annual International Conference of the IEEE, 2008.

[10] Raventós, A. and Zaidi, M., “Automating Neurological Disease Diagnosis Using Structural MR Brain Scan Features.”.

[11] Yann LeCun, Léon Bottou, Yoshua Bengio, and Patrick Haffner, “Gradient-based learning applied to document recognition,” Proceedings of the IEEE, vol. 86, no. 11, pp. 2278–2324, 1998.

[12] Yangqing Jia, Evan Shelhamer, Jeff Donahue, Sergey Karayev, Jonathan Long, Ross Girshick, Sergio Guadarrama, and Trevor Darrell., “Caffe: Convolutional architecture for fast feature embedding,” In Proceedings of the ACM International Conference on Multimedia, pp. 675–678, 2014.

[13] Jiquan Ngiam, Aditya Khosla, Mingyu Kim, Juhan Nam, Honglak Lee, and Andrew Y. Ng, “Multimodal deep learning,” in In Proceedings of the 28th international conference on machine learning (ICML-11), 2011.

[14] Dumitru Erhan, Yoshua Bengio, Aaron Courville, Pierre-Antoine Manzagol, Pascal Vincent, and Samy Bengio, “Why does unsupervised pre-training help deep learning?,” Journal of Machine Learning Research, vol. 11, pp. 625–660, 2010.

[15] J. Schmidhuber, “Deep learning in neural networks: An overview,” Neural Networks, vol. 61, pp. 85–117, 2015.

[16] Itamar Arel, Derek C. Rose, and Thomas P. Karnowski, “Deep machine learning-a new frontier in artificial intelligence research [research frontier],” Computational Intelligence Magazine, IEEE, vol. 5, no. 4, pp. 13–18, 2010.

[17] Christian Szegedy, Wei Liu, Yangqing Jia, Pierre Sermanet, Scott Reed, Dragomir Anguelov, Dumitru Erhan, Vincent Vanhoucke, and Andrew Rabinovich, “Going deeper with convolutions.,” In Proceedings of the IEEE Conference on Computer Vision and Pattern Recognition, pp. 1–9, 2015.

[18] Limin Wang, Zhe Wang, Wenbin Du, and Yu Qiao, “Object-scene convolutional neural networks for event recognition in images,” in In Proceedings of the IEEE Conference on Computer Vision and Pattern Recognition Workshops, 2015.

[19] Alex Krizhevsky, Ilya Sutskever, and Geoffrey E. Hinton, “Imagenet classification with deep convolutional neural networks,” In Advances in neural information processing systems, pp. 1097–1105, 2012.

[20] D. G. Lowe, “Distinctive image features from scale-invariant keypoints,” International journal of computer vision, vol. 60, pp. 91–110, 2004.

[21] Karen Simonyan, Andrew Zisserman, “Very deep convolutional networks for large-scale image recognition,” arXiv preprint arXiv 1409.1556, 2014.

[22] Kaiming He, Xiangyu Zhang, Shaoqing Ren, and Jian Sun, “Deep residual learning for image recognition,” arXiv preprint arXiv:1512.03385, 2015.

[23] Clifford R. Jack, Matt A. Bernstein, Nick C. Fox, Paul Thompson, Gene Alexander, Danielle Harvey, Bret Borowski et al., “The Alzheimer’s disease neuroimaging initiative (ADNI): MRI methods,” Journal of Magnetic Resonance Imaging, vol. 27, no. 4, pp. 685–691, 2008.

[24] Suk, Heung-Il, and Dinggang Shen, “Deep learning-based feature representation for AD/MCI classification,” In International Conference on Medical Image Computing and Computer-Assisted Intervention, pp. 583–590, 2013.

[25] Heung-Il Suk, Seong-Whan Lee, Dinggang Shen, “Latent feature representation with stacked auto-encoder for AD/MCI diagnosis,” Brain Structure and Function, vol. 220, no. 2, pp. 841–859, 2015.

[26] Heung-Il Suk, Dinggang Shen, “Deep learning in diagnosis of brain disorders,” In Recent Progress in Brain and Cognitive Engineering, pp. 203–213, 2015.

[27] Suk, Heung-Il, Seong-Whan Lee, Dinggang Shen, “Hierarchical feature representation and multimodal fusion with deep learning for AD/MCI diagnosis,” NeuroImage, vol. 101, pp. 569–582, 2014.

[28] Adrien Payan, Giovanni Montana, “Predicting Alzheimer’s disease: a neuroimaging study with 3D convolutional neural networks,” arXiv preprint arXiv:1502.02506, p. 2015.

[29] Siqi Liu, Sidong Liu, Weidong Cai, Hangyu Che, Sonia Pujol, Ron Kikinis, Dagan Feng, and Michael J. Fulham, “Multimodal Neuroimaging Feature Learning for Multiclass Diagnosis of Alzheimer’s Disease,” IEEE Transactions on Biomedical Engineering, vol. 62, no. 4, pp. 1132–1140, 2015.

[30] E. Arvesen, “Automatic Classification of Alzheimer’s Disease from Structural MRI,” 2015.

[31] Fayao Liu, and Chunhua Shen, “Learning Deep Convolutional Features for MRI Based Alzheimer’s Disease Classification,” arXiv preprint arXiv:1404.3366, 2014.

[32] Siqi Liu, Sidong Liu, Weidong Cai, Hangyu Che, Sonia Pujol, Ron Kikinis, Michael Fulham, and Dagan Feng, “High-level feature based PET image retrieval with deep learning architecture,” Journal of Nuclear Medicine, vol. 1, pp. 2028–2028, 2014.

[33] Tom Brosch, Roger Tam, “Manifold learning of brain MRIs by deep learning,” in In International Conference on Medical Image Computing and Computer-Assisted Intervention, Berlin, 2013.

[34] Ladislav Rampasek, Anna Goldenberg, “TensorFlow: Biology’s Gateway to Deep Learning?,” Cell systems, vol. 2, no. 1, pp. 12–14, 2016.

[35] Alexander de Brebisson, Giovanni Montana, “Deep neural networks for anatomical brain segmentation,” in In Proceedings of the IEEE Conference on Computer Vision and Pattern Recognition Workshops, Boston, 2015.

[36] Earnest Paul Ijjina, and Chalavadi Krishna Mohan, “Hybrid deep neural network model for human action recognition,” Applied Soft Computing, 2015.

[37] Xiaofeng Zhu, Heung-Il Suk, Dinggang Shen, “A novel matrix-similarity based loss function for joint regression and classification in AD diagnosis,” NeuroImage, vol. 100, pp. 91–105, 2014.

[38] Chen Zu, Biao Jie, Mingxia Liu, Songcan Chen, Dinggang Shen, Daoqiang Zhang, Alzheimer’S Disease Neuroimaging Initiative, “Label-aligned multi-task feature learning for multimodal classification of Alzheimer’s disease and mild cognitive impairment,” Brain imaging and behavior, pp. 1–12, 2015.

[39] Siqi Liu, Sidong Liu, Weidong Cai, Hangyu Che, Sonia Pujol, Ron Kikinis, Dagan Feng, Michael J. Fulham, “Multimodal Neuroimaging Feature Learning for Multiclass Diagnosis of Alzheimer’s Disease,” IEEE Transactions on Biomedical Engineering, vol. 62, no. 4, pp. 1132–1140, 2015.

[40] M. Liu, D. Zhang, E. Adeli-Mosabbeb, D. Shen, “Inherent Structure Based Multi-view Learning with Multi-template Feature Representation for Alzheimer’s Disease Diagnosis.,” IEEE transactions on bio-medical engineering, 2015.

[41] Feng Li, Loc Tran, Kim-Han Thung, Shuiwang Ji, Dinggang Shen, Jiang Li, “A robust deep model for improved classification of AD/MCI patients,” IEEE journal of biomedical and health informatics, vol. 19, no. 5, pp. 1610–1616, 2015.

[42] Ehsan Hosseini-Asl, Robert Keynton, Ayman El-Baz., “Alzheimer’s disease diagnostics by adaptation of 3D convolutional network,” in In Image Processing (ICIP), 2016 IEEE International Conference on, 2016.

[43] S. Smith., “Fast robust automated brain extraction,” Human Brain Mapping, vol. 17(3), no. November 2002, pp. 143–155, 2002.

[44] Jenkinson, M., Bannister, P., Brady, J. M. and Smith, S. M., “Improved Optimisation for the Robust and Accurate Linear Registration and Motion Correction of Brain Images,” NeuroImage, vol. 17(2), pp. 825–841, 2002.

[45] Gwenaëlle Douaud, Stephen Smith, Mark Jenkinson, Timothy Behrens, Heidi Johansen-Berg, John Vickers, Susan James et al., “Anatomically related grey and white matter abnormalities in adolescent-onset schizophrenia,” Brain, vol. 130, no. 9, pp. 2375–2386, 2007.

[46] Saman Sarraf, and Ghassem Tofighi, “Classification of Alzheimer’s Disease using fMRI Data and Deep Learning Convolutional Neural Networks,” arXiv preprint arXiv:1603.08631, 2016.

